# Probabilistic ecological risk assessment for deep-sea mining: a Bayesian Network for Chatham Rise, SW Pacific Ocean

**DOI:** 10.1101/2023.11.28.569078

**Authors:** Laura Kaikkonen, Malcolm R. Clark, Daniel Leduc, Scott D. Nodder, Ashley A. Rowden, David A. Bowden, Jennifer Beaumont, Vonda Cummings

## Abstract

Increasing interest in seabed resource use in the ocean is introducing new pressures on deep-sea environments, the ecological impacts of which need to be evaluated carefully. The complexity of these ecosystems and the dearth of comprehensive data pose significant challenges to predicting potential impacts. In this study, we demonstrate the use of Bayesian Networks (BNs) as a modelling framework to address these challenges and enhance the development of robust quantitative predictions concerning the effects of human activities on deep-seafloor ecosystems. The approach consists of iterative model building with experts, and quantitative probability estimates of the relative decrease in abundance of different functional groups of benthos following seabed mining. The model is then used to evaluate two alternative seabed mining scenarios to identify the major sources of uncertainty associated with the mining impacts. By establishing causal connections between the pressures associated with potential mining activities and various components of the benthic ecosystem, our model offers an improved comprehension of potential impacts on the seafloor environment. We illustrate this approach using the example of potential phosphorite nodule mining on the Chatham Rise, offshore Aotearoa/New Zealand, SW Pacific Ocean, and examine ways to incorporate knowledge from both empirical data and expert assessments into quantitative risk assessments. We further discuss how ecological risk assessments can be constructed to better inform decision-making, using metrics relevant to both ecology and policy. The findings from this study highlight the valuable insights that BNs can provide in evaluating the potential impacts of human activities. However, continued research and data collection are crucial for refining and ground truthing these models and improving our understanding of the long-term consequences of deep-sea mining and other anthropogenic activities on marine ecosystems. By leveraging such tools, policymakers, researchers, and stakeholders can work together towards human activities in the deep sea that minimise ecological harm and ensure the conservation of these environments.

## 1 Introduction

Interest in seabed mining, deep-sea fishing, and oil and gas exploration is increasing in the deep sea (Jouffray et al., 2020; Ramirez-Llodra et al., 2011). In order to effectively manage these activities, predicting their impacts on deep-sea environments and ecosystems prior to resource consent approval is crucial. However, the complexity of deep-sea ecosystems and the lack of comprehensive data often make predicting impacts challenging (Smith et al., 2020). It is therefore important to conduct robust environmental risk assessments (ERAs) that take into account the potential risks and uncertainties associated with activities that can impact deep-sea habitats and communities.

Significant mineral resources have been identified in various parts of the global ocean, including areas of the Pacific, Atlantic, and Indian oceans (Ellis et al., 2017; Miller et al., 2018). Seabed mining activities are still in the exploration and development phase and no commercial mining operations have yet taken place, but concerns have been raised over their potential to harm deep-sea ecosystems (van Dover et al., 2017). Deep-sea mining is expected to have an impact on all levels of marine ecosystems, from the water column to the seafloor (Drazen et al., 2020; Miljutin et al., 2011; Orcutt et al., 2020). Many studies have examined the potential environmental consequences of mining by using field studies, laboratory experiments, and modelling (reviewed by Jones et al., (2017). However, despite the valuable insights provided by these studies, there is only a partial understanding of the environmental impacts of deep-sea mining. Current knowledge gaps include uncertainties about the scale and duration of the effects, the potential for cumulative impacts over time, and the extent and speed of ecosystem recovery following disturbance (Amon et al., 2022). Furthermore, it is unclear to what extent the disturbance studies conducted in a small area or in a laboratory can be scaled up to industrial mining operations (Clark, Durden, and Christiansen 2020). Limited baseline data and the difficulty of access to some of the remote areas where deep-sea mining is proposed make it challenging to accurately assess the environmental risks associated with such mining activities (Smith et al., 2020).

ERAs are an important tool to help environmental managers evaluate the risks associated with mining operations. In the deep sea, ERAs can be particularly useful to support decision-making, due to limitations of baseline data and of information on ecosystem responses to external disturbances. However, most current ERAs estimate risk based upon the vulnerability of the environment through semi-quantitative scoring (Boschen-Rose et al., 2021; Washburn et al., 2019), offering an overview of the risks without quantitative estimates of the actual ecosystem impacts. To account for the uncertainties related to such lack of data, probabilistic modelling has been increasingly used in ERAs (Kaikkonen et al., 2021).

Bayesian networks (BNs) are graphical probabilistic models that provide an alternative to commonly used scoring procedures in ERAs (Kelly et al., 2013; Pearl 2010). In a risk assessment context, BNs illustrate the modelled system as a network of causal influences. BNs are composed of a directed acyclic graph (the structure of the network) with quantitative connections between the variables (or nodes). The strength of each connection between variables is described through conditional probabilities (Pearl 1986), thus representing a joint probability distribution over a set of variables. The dependencies between variables propagate through the network and influence the probabilities of other nodes and may be updated as new information about the nodes becomes available. This facet of the model enables the integration of new data or evidence in the model, and the network can be queried under different scenarios to calculate the posterior probability of all other nodes within the BN (Kelly et al., 2013; Pearl 2010).

Unlike traditional scoring procedures, BNs allow for the estimation of not only the most likely outcome but also the uncertainty associated with the estimates by providing a probability distribution over all the possible values of each variable (Fenton and Neil 2012; Nielsen and Jensen 2009). BNs can synthesise outcomes of multiple scenarios and accommodate inputs from multiple sources, including simulations, empirical data, and expert knowledge (e.g., Bulmer et al., 2022; Wade et al., 2021), making them well-suited for data-poor cases. Additionally, given their modular structure, BNs can support integrative modelling combining different submodels, such as management decision networks (Marcot and Penman 2019).

In this paper, we apply BNs in a case study focused on potential phosphate nodule mining on the Chatham Rise, offshore Aotearoa/New Zealand, SW Pacific Ocean. Drawing on a combination of field observations, laboratory experiments, and expert knowledge, we estimate the likelihoods of impacts on benthic fauna under a high disturbance and an intermediate disturbance seabed mining scenarios.

## 2 Material and methods

### 2.1 Chatham Rise phosphate nodule mining case study

#### 2.1.1 Background

In 2013, a New Zealand company, Chatham Rock Phosphate (CRP), applied for and was granted a Minerals Mining Permit by the New Zealand Government for phosphate nodule extraction from the seafloor on the Chatham Rise, located in the central eastern region of New Zealand’s 200 nautical mile Exclusive Economic Zone (EEZ) (Fig.1). The depth of the crest of the Chatham Rise is 200 to 500 m and its flanks deepen to more than 2000 m to the north and south (Nodder et al., 2012). The area is characterised by high primary productivity, with dynamic oceanographic conditions characterised by variable currents and interweaving water masses associated with the Subtropical Frontal Zone (Collins et al., 2023; Safi et al., 2023). The sediments covering the crest are predominantly organic-rich, glauconitic muddy sands and sandy muds, with phosphorite nodules and hardgrounds on top and within the sediment (Cullen 1987; Nelson et al., 2022; Norris 1964). The seafloor communities in the area are characterised by a wide range of invertebrate species, many of which are infaunal, although some species occupy habitat niches either on top of the sediment (epifauna) or just above the seafloor within the hyperbenthos (Compton et al., 2013). Corals and other sessile epifaunal organisms, such as sponges, also live attached to hard substrates such as phosphorite nodules or rock outcrops (Dawson 1984; Leduc et al., 2015; Nodder et al., 2012; Rowden et al., 2014).

**Figure 1.**
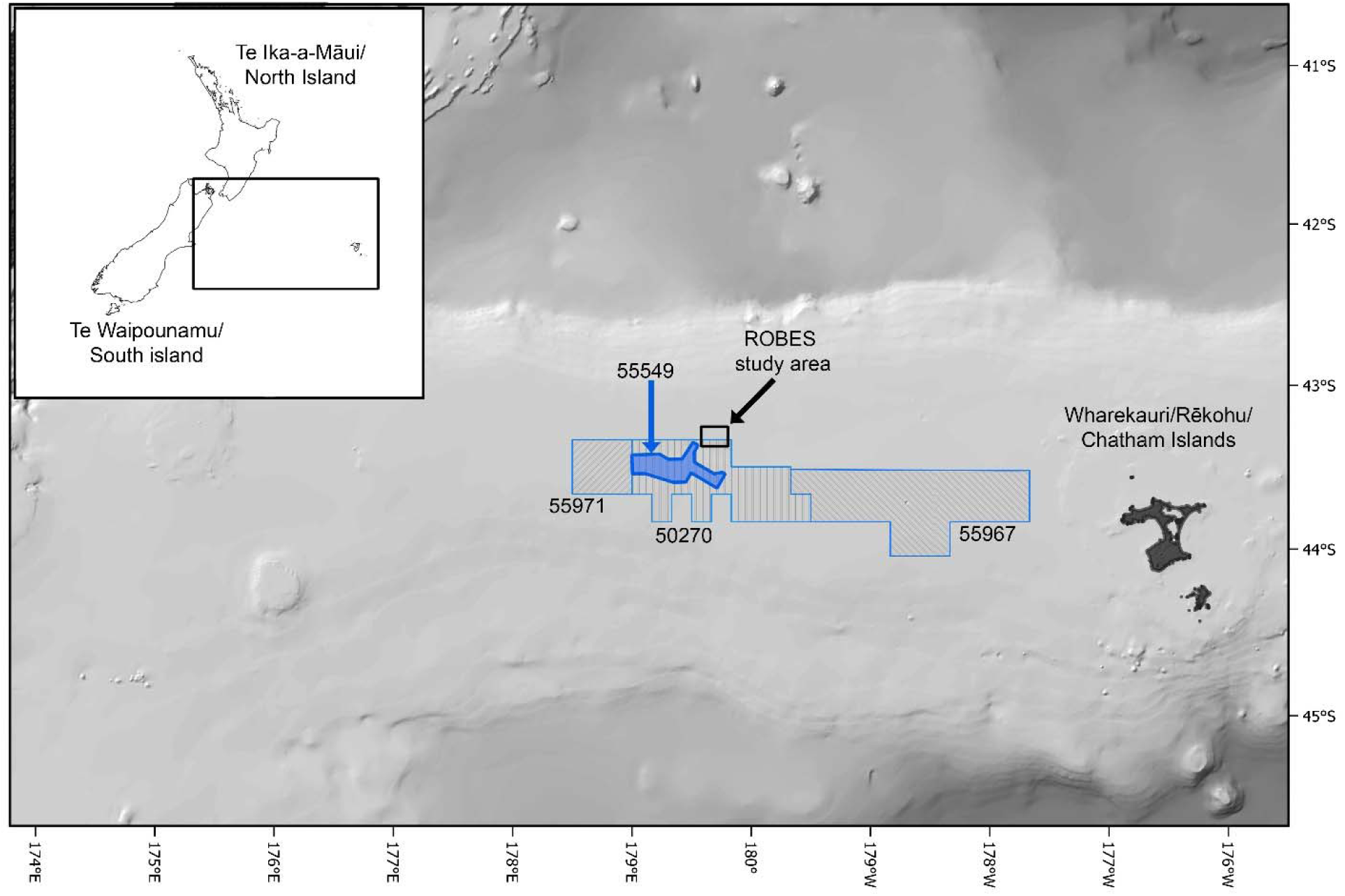
Map of study area on the Chatham Rise, offshore of New Zealand. The numbered polygons denote the CRP Minerals Mining Permit (55549) and previous Mineral Prospecting Permit areas.

The proposed mining operation was to extract phosphorite nodules from the seafloor using a trailing suction drag-head and to mechanically process the nodules on board the mining support vessel. Nodules larger than 2 mm in diameter would be separated from other sediment material and the waste would then be discharged close to the seafloor via a discharge pipe (Chatham Rock Phosphate 2014). The mining would be carried out over separate mining blocks, each covering an area of 5 km by 2 km and taking approximately 14 weeks to complete mining operations. However, in 2015, the marine consent application to carry out the mining of phosphorite nodules was denied, due in part to uncertainty surrounding the potentially adverse effects on biological communities, including impacts caused by suspended and deposited sediment (NZ EPA 2015).

In order to address the scientific uncertainties related to impacts from seabed sediment disturbance, the “Resilience of deep-sea benthic communities to the effects of sedimentation” (ROBES) programme gathered information on various environmental factors and benthic fauna. The programme consisted of both field and laboratory simulations to characterise the benthic effects of an artificial physical seabed disturbance and associated sediment plume on the Chatham Rise crest in the vicinity of the proposed CRP phosphorite mining area (Fig. 1). The experimental seabed disturbances, although not equivalent to actual seabed mining, were anticipated to provide important insights into the impacts of deep-sea mining and other significant benthic disturbances such as bottom trawling (Clark et al., 2018).

#### 2.1.2 Data

The data used in this study originate from the field and laboratory measurements collected through the ROBES programme in 2018–2020. The fieldwork took place on the northern edge of the Chatham Rise crest at depths of 400–500 m (Fig. 1). The study area was surveyed in 2018 and 2019, then artificially disturbed using a mechanical disturber and sampled immediately after the disturbance in 2019 and one year later in June 2020 (Clark et al., 2021). A diverse range of data was collected to characterise the site, encompassing oceanographic (Acoustic Doppler current profiler (ADCP), ocean glider, moorings, conductivity, temperature, and depth (CTD), acoustics) and nearbed sediment conditions (benthic landers, sediment trap moorings, multicorer, onboard sediment experiments), and benthic communities. Environmental baseline conditions were determined using year-long ADCP moorings and glider and CTD deployments during the ROBES voyages, followed by modelling and analysis of satellite remote-sensing data (Collins et al., 2023). Information on baseline nearbed particle and organic carbon fluxes were derived from bi-weekly and daily sampling using moored and lander sediment traps, respectively. Macroinfauna and meiofauna samples were collected before and after the disturbance using a multicorer with replicate samples from sites that were directly impacted by the mechanical disturber and from near-field areas that were expected to be subject to sedimentation (see Clark et al., 2019; Clark et al., 2021 for details of the sampling protocol). In addition to the field sampling, live sponges and corals were transported back to the laboratory to assess their response to different concentrations and frequencies of suspended sediment over time (for details, see Mobilia et al., 2021, 2023).

### 2.2 BN modelling

#### 2.2.1 Model development and variable selection

A conceptual influence diagram synthesising the impacts of deep-sea mining on the Chatham Rise was developed in a series of workshops with experts (Appendix 1, Table S2). Given the complexity of the Chatham Rise ecosystem we focus only on benthic ecosystem impacts associated with seabed mining. Due to lack of empirical data, we did not consider impacts from noise and vibrations associated with the mining activities in our assessment nor the environmental effects on certain components of the marine foodweb, such as bacteria, marine mammals, and seabirds.

The causal network resulting from the expert elicitation was developed into a BN model in an iterative manner by selecting key variables to evaluate and defining relevant variable states. To facilitate model quantification and ensure model parsimony, the number of parent nodes was limited whenever possible (Chen and Pollino, 2012).

Discrete variable states were defined based on data, literature, and expert views (Table 1, full references in Appendix 1, Table S3). Variable states were chosen to represent likely variations in the variables of interest in the context of seabed disturbance on the Chatham Rise. For variables that describe the implementation of the potential mining activity (hereafter ‘operational variables’), variable states were drawn from the environmental consent application prepared by CRP (Chatham Rock Phosphate, 2014). The states of the physicochemical variables, describing the environmental conditions and associated changes from mining, were defined by experts based on field observations, expert knowledge, and primary literature. For variables that directly affect benthic fauna, such as suspended sediment concentration, the states were set to reflect biologically relevant thresholds whenever possible (e.g., Hewitt and Lohrer, 2013; Mobilia et al., 2021, 2023). As a result, the states do not always follow a continuous scale nor cover all possible values of a variable but were selected to represent likely outcomes from different disturbance events (see Appendix1 Table S3 for rationale for variable discretisation).

**Table 1.** Physical and environmental model variables and the methods used to discretise and parameterise the variables used in the Bayesian Network modelling. Random variables refer to variables with an associated probability distribution, whereas decision variables describe processes that are assumed to be controlled by the party responsible for the extraction activity and are thus non-random. Full rationale supporting the variable states and parameterisation with references are contained in Appendix 1.

| **Variable name** | **Description** | **Variable type** | **Variable states** | **Discretisation based on** | **Parameterisation based on** |
| --- | --- | --- | --- | --- | --- |
| Depth of extracted sediment | Depth of sediment extracted by the mining tool | Decision variable | <10cm / 10-30 cm / >30cm | Literature, expert opinion | Not applicable |
| Processing return technique | Depth of processing return water and sediment | Decision variable | 10m from seafloor / at the seafloor | Literature | Not applicable |
| Mining intensity | Proportion of area mined within a discrete mining block | Decision variable | 50% / 75% / 100% | Expert opinion | Not applicable |
| Distance from the mining block | Distance from the mining block | Decision variable | Inside mining block / Near-field / Far-field | Literature, expert opinion | Not applicable (decision variable) |
| Volume of extracted sediment | Volume of sediment removed by a mining operation tool (as the initial removal) | Random variable | Low/ Medium /High | Literature, expert opinion | Literature, expert assessment |
| Particle size composition of the extracted sediment | Proportion of fine and coarser particles of the extracted substrate and in the composite sediment plume | Random variable | Mostly silt and clay (fine) / mix of fine and coarse particles / mostly coarse (sand and gravel) | Data, expert opinion | Field surveys |
| Sediment contaminants | Concentration of potentially harmful substances in the sediment and the mineral material to be extracted | Random variable | Low/ Medium /High | Literature, expert opinion | Literature, expert assessment |
| Nodule removal | Proportion of phosphorite nodules removed from a discrete mining block. | Decision variable | Yes / No | Not applicable | Not applicable (decision variable) |
| Suspended sediment | Total suspended solids concentration near the seafloor resulting both from the processing return and mining tool operation | Random variable | Low: <10 mg L<sup>-1</sup><br>/ Moderate: 11-50 mg L<sup>-1</sup><br>/ High:>100 mg L<sup>-1</sup> | Data, literature | Expert assessment informed by literature |
| Sediment deposition | Depth of sediment deposited from the composite suspended sediment plume | Random variable | Low: <1-2 cm /<br>Moderate: 2-5 cm /<br>High:10-25cm | Data, literature | Field measurements and expert assessment |
| Contaminant release | Release of contaminants from the sediment plume to the seabed water column | Random variable | Nonsignificant /<br>Significant release of contaminants | Data, literature | Expert assessment |

| Changes in sediment characteristics | Measure of changes in the sediment environment affecting habitat quality for benthic organisms. | Random variable | Minor to no changes / Significant changes | Data, expert opinion | Field measurements and expert assessment |
| --- | --- | --- | --- | --- | --- |

As the number of species found in the study area is too high to assess the impacts on each species or taxon separately, we reduced this complexity by grouping organisms into functional groups (Bremner 2008). The functional groups were assigned based on traits that have been shown to influence the organisms’ response to seafloor disturbance and recovery potential (e.g., body size, feeding habit, position in sediment, and mobility; Hewitt et al., 2018), using previously created groupings for the Chatham Rise as a starting point (Lundquist et al., 2018). These trait-based groups encompassed a range of faunal groups across meiofauna, macroinfauna, epibenthos and hyperbenthos (Table 2).

**Table 2.** Biological model variables describing the benthic faunal functional groups and the methods used to discretise and parameterise the variables used in the Bayesian Network modelling. Random variables refer to variables with an associated probability distribution. Full rationale supporting the variable states and parameterisation with references are contained in the conditional probability tables in Appendix 1.

| Variable name and abbreviation used in the paper | Example taxa | Variable type | Variable states | Discretisation method | Parameterisation method |
| --- | --- | --- | --- | --- | --- |
| Sessile encrusting suspension feeders (SCSF) | Encrusting bryozoans, corals | Random variable | % decrease in abundance | Expert opinion | Expert assessment |
| Sessile encrusting filter-feeders (SCFF) | Encrusting sponges | Random variable | % decrease in abundance | Expert opinion | Expert assessment |

| **Variable name and abbreviation used in the paper** | **Example taxa** | **Variable type** | **Variable states** | **Discretisation method** | **Parameterisation method** |
| --- | --- | --- | --- | --- | --- |
| Sessile erect suspension feeders (SESF) | Branching bryozoans, crinoids and corals (stony and octocorals) | Random variable | % decrease in abundance | Expert opinion | Laboratory experiment and expert assessment |
| Sessile erect filter-feeders (SEFF) | Erect sponges | Random variable | % decrease in abundance | Expert opinion | Laboratory experiment and expert assessment |
| Soft-bodied erect suspension and filter feeders (SSBM) | Small soft bodied hydroids, ascidians, small bryozoans | Random variable | % decrease in abundance | Expert opinion | Expert assessment |
| Deep meiofauna (DM) | Mainly nematodes | Random variable | % decrease in abundance | Expert opinion | Field measurements and expert assessment |
| Surface meiofauna (SM) | Mixed community of meiofauna | Random variable | % decrease in abundance | Expert opinion | Field measurements and expert assessment |
| Small sessile infauna (SSI) | Paraonid and capitellid polychaetes, small bivalves | Random variable | % decrease in abundance | Expert opinion | Field measurements and expert assessment |
| Small mobile infauna (SMI) | Mobile deposit feeders and small scavengers (amphipods, small crustaceans) | Random variable | % decrease in abundance | Expert opinion | Field measurements and expert assessment |
| Large sessile infauna (LSI) | Tube-forming polychaetes, tube-building amphipods, large bivalves | Random variable | % decrease in abundance | Expert opinion | Field measurements and expert assessment |
| Large mobile macroinfauna (LMM) | Large burrowing polychaetes, some arthropods, large bivalves, asteroids | Random variable | % decrease in abundance | Expert opinion | Field measurements and expert assessment |
| Mobile deposit feeding or grazing epibenthos (MGE) | Mobile deposit feeders, surface dwelling species, e.g., spatangoid echinoids, holothurians, ophiuroids, gastropods | Random variable | % decrease in abundance | Expert opinion | Expert assessment |
| Mobile predatory or scavenging epibenthos (MPE) | Mobile surface crawling predators & scavengers, e.g., squat lobsters, crab, scampi, gastropods | Random variable | % decrease in abundance | Expert opinion | Expert assessment |
| Predatory or Scavenging hyperbenthos (PH) | Small swimming crustaceans (mysids, amphipods) | Random variable | % decrease in abundance | Expert opinion | Expert assessment |
| Grazing or deposit-feeding hyperbenthos (GH) | Holothurians, gastropods, others | Random variable | % decrease in abundance | Expert opinion | Expert assessment |

#### 2.2.2 Model parameterisation

Within a BN, the magnitudes of impacts are illustrated through conditional dependencies. The probabilities of each value of the ‘child’ node, conditioned on every possible combination of values of the ‘parent’ nodes, can be drawn from data, expert opinion, or a combination of these two inputs (Barber 2012). Conditional probabilities (summarised in conditional probability tables, CPTs) were derived from a combination of data and expert assessment (Tables 1-2). Where data on the impacts were available, we used experimental and field data to inform the probability distributions and complemented these with expert assessment. As BNs require probability estimates for all the combinations of variable states, the configurations of parent variables that were not applied in the field study were estimated by experts through the following procedure.

In order to generalise the impacts of stressors on the different functional groups and to reduce the elicitation burden on experts, we applied an interpolation method (Barons et al., 2022) to derive missing probability distributions. The method assumes that the child node can be estimated through a beta distribution and requires the user to identify both a “best case” and a “worst case” distribution for the child node. In this context, “best” and “worst” pertain to distributions where the parents affecting the node are at their highest and lowest points for the child, respectively. The probability distributions for all other combinations of parent variables are then inferred by interpolating between these extremes. This procedure is accomplished using a series of weights assigned to the parent states, which are employed to interpolate between the parameters of a beta distribution.

Where data on the impacts were available, empirical data were used as the starting point which corresponded to the best case scenario, and experts were then asked to estimate: 1) the relative importance of each of the stressors on each functional group, and 2) the probability distribution for the relative decrease in abundance (compared to pre-mining abundance) of each functional group under the worst case scenario (i.e., each stressor at the maximum state). The interpolation method can be applied to variables with ranked ordinal states (e.g., low to high). For nodes where such ordinal ranking was not appropriate, we used the Application for Conditional probability Elicitation (ACE; Hassall et al., 2019) to initialise the CPTs, which were then reviewed with experts. All CPTs, as well as a more detailed description of the elicitation protocol, are available at https://github.com/lkaikkonen/CR-ERA, including comments on the rationale underlying the probability distributions for each response variable.

All CPTs were reviewed with the experts and adjusted when deemed necessary to ensure consistency in the estimated impacts. In addition to the probability estimates, experts evaluated their confidence in the estimates for each of the conditional probability tables. The resulting CPTs were incorporated in the BN model created in R software. The modelling was conducted using R 3.6.3, with the R package *bnlearn* (Scutari 2009). Full details of the model with the R scripts and the conditional probability tables are available at https://github.com/lkaikkonen/CR-ERA.

#### 2.2.3 Modelling framework and model structure

We consider three spatial domains in the model: the area inside the mining block that is mined (inside), areas immediately adjacent to the mining block that are not mined (near-field), and areas further from the mining area that are still expected to be within the zone impacted by the mining activities (far-field) (Fig. 2). Unimpacted areas beyond far-field are not included in the model. In the CRP mining proposal, the mining block covers an area of 5 km by 2 km. For the purposes of this study, we conceptualise the near-field area to extend approximately 0.5 km from the mined area and the far-field to extend to 5 km from the mined area. However, it is important to note that the size of the near-field and far-field are relative to the scale of disturbance (usually defined by the extent of detectable impacts) and will therefore vary for other types of disturbances and areas. As our model is not spatially explicit, we assume homogeneous impacts inside all spatial domains.

**Figure 2.**
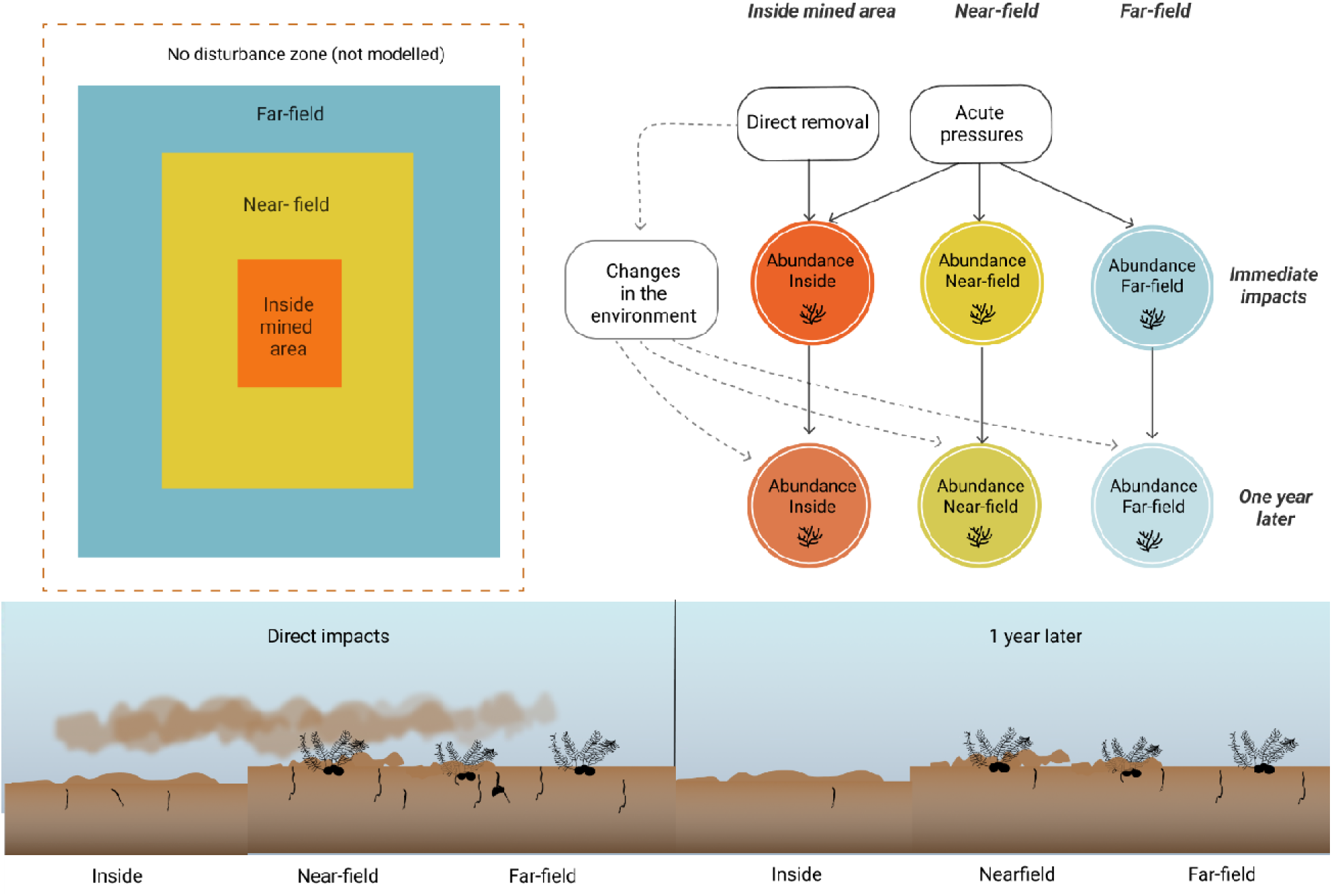
General modelling framework for impacts of seabed mining-related disturbance in time and space for any given functional group in the BN model. By defining the effects for the separate time steps, the impacts may be assessed jointly or separately for each discrete area (depicted by the squares, upper left panel). The lower panel illustrates the spatial and temporal distribution of the pressures. Immediate impacts consist of direct extraction of sediment and nodules within the mining block, and elevated suspended sediment concentrations and redeposition of sediment in all areas. Impacts after one year are a result of the altered sedimentary environment which affects the recovery potential of benthic organisms. The magnitude and likelihood of all these changes are variable and included in the probability assessments (see text for details).

We estimated the impact on benthic fauna as a decrease in abundance relative to the pre-disturbed state. For most faunal groups this decrease in abundance translates to mortality, but as our model also includes mobile fauna that may leave the area but are not killed by the disturbance, we use the term ‘decrease in abundance’ throughout the paper. As a simplification, we only assess decrease in abundance, although some faunal groups may temporarily increase in abundance after seabed disturbance (e.g., Bigham et al., 2023; Pranovi et al., 2000). We divided the abundance variables into two time-steps: immediately after disturbance and one year after disturbance based on the available data from the ROBES experiments (Fig. 2). We separated the decrease in abundance immediately after mining into direct and indirect decrease in abundance (see Appendix 1 for full description of the modelling framework). Within this framework, some of the organisms will be removed during the direct extraction process (direct impact), depending on the mining efficiency and depth. The remaining fauna will be exposed to indirect impacts (in our model these are sedimentation and impacts from toxic substance release) that will describe the acute impacts on them (details in Appendix 1, following Kaikkonen et al., 2021). Recovery is assessed separately, and any changes in the seafloor environment (e.g., sediment composition, nodule removal) only affect the recovery of organisms, not immediate changes in abundance. Any subsequent time steps will depend on the faunal abundance at the previous time step, recovery potential of the functional group, and changes in the habitat quality (such as changes in sediment characteristics and food availability). The division of direct and indirect pressures that affect the decrease in abundance also allows us to evaluate the impacts of mining in both the mined and unmined areas within the same model.

### 2.3 Application: Disturbance scenarios and use of the BN model

BNs enable evaluation of various scenarios and computation of posterior probabilities based on new knowledge (Pearl 1986). Through BNs, operational parameters can be modified to analyse the effects of different types of seabed mining (or other types of seabed disturbance) and their impact on benthos. The joint probability distribution in the BN can be used to query the effects of multiple pressures on specific ecosystem components, assess associated risks, and identify the variables that should be monitored for an improved understanding of the impacts (Carriger et al., 2016).

In order to assess how changes in the magnitude of disturbance affect benthic fauna, we queried the network on two alternative mining scenarios. These scenarios, which we define as a combination of specific states of the decision variables that describe the overall mining process, are assumed to be controlled by the mining operator (Table 3). In the first scenario, hereafter ‘High disturbance’, the entire mined area (Fig. 2) was disturbed, and sediments were disrupted to deeper than 30 cm. For the second scenario, hereafter ‘Intermediate disturbance’, 50% of the mined area was disturbed, and sediments were disrupted to less than 10 cm depth. The High disturbance scenario was defined by experts based on the description of a proposed mining operation for phosphorite nodules on the Chatham Rise (Chatham Rock Phosphate 2014), while the Intermediate disturbance scenario was based on the anticipated disturbance from other types of mining operations (e.g., surface nodule extraction as proposed for polymetallic nodule mining, e.g., Muñoz-Royo et al., 2022) and could also be applied to describe low-penetration bottom trawling (Eigaard et al., 2016). All the other variables in the model are further affected by these decision variables. Note that the model can be queried for any combination of variables, we have presented only a limited number of possible outcomes.

**Table 3.** States of the operational variables associated with the two seabed mining disturbance scenarios.

| Scenario | Operational variables |  |  |  |
| --- | --- | --- | --- | --- |
|  | Area disturbed inside the mining block | Depth of sediment disturbance | Plume release technique | Description |
| High disturbance | 100% | >30 cm | At the seafloor | High impact seabed mining operations |
| Intermediate disturbance | 50% | <10 cm | At the seafloor | Surface collector operations |

Developing models in data-limited settings presents a challenge for validating these models using conventional statistical methods. This difficulty arises from the impossibility of testing the model against an independent dataset that was not employed during the model’s development and quantification process. In addition, conventional sensitivity analyses do not provide much insight as the model structure has been defined by experts. Therefore, the BN was qualitatively evaluated in a series of meetings attended by experts, during which the model and its outcomes were presented, discussed, and agreed upon. A full overview of the model building, and application is given in Figure 3.

**Figure 3.**
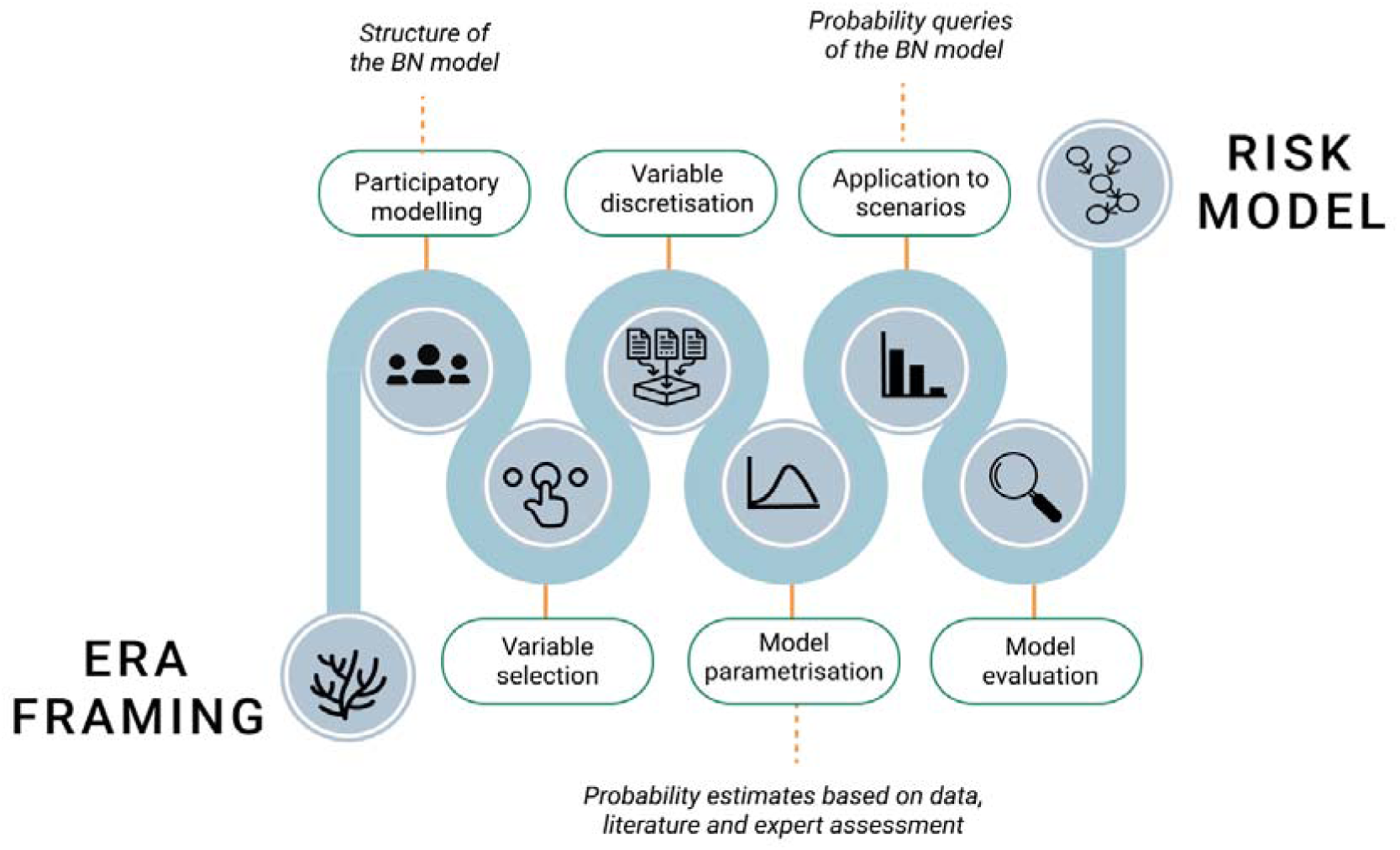
Overview of modelling process. ERA = environmental risk assessment.

## 3 Results

The causal mapping and model building process resulted in a BN model for the Chatham Rise with 73 variables and 154 connections (Fig. 4). The model has seven independent variables describing the two disturbance scenarios and environmental conditions/pressures caused by mining that further cascade down to responses in benthic fauna. In this section we present results on the joint probabilities queried on the two disturbance scenarios for the Chatham Rise environment.

**Figure 4.**
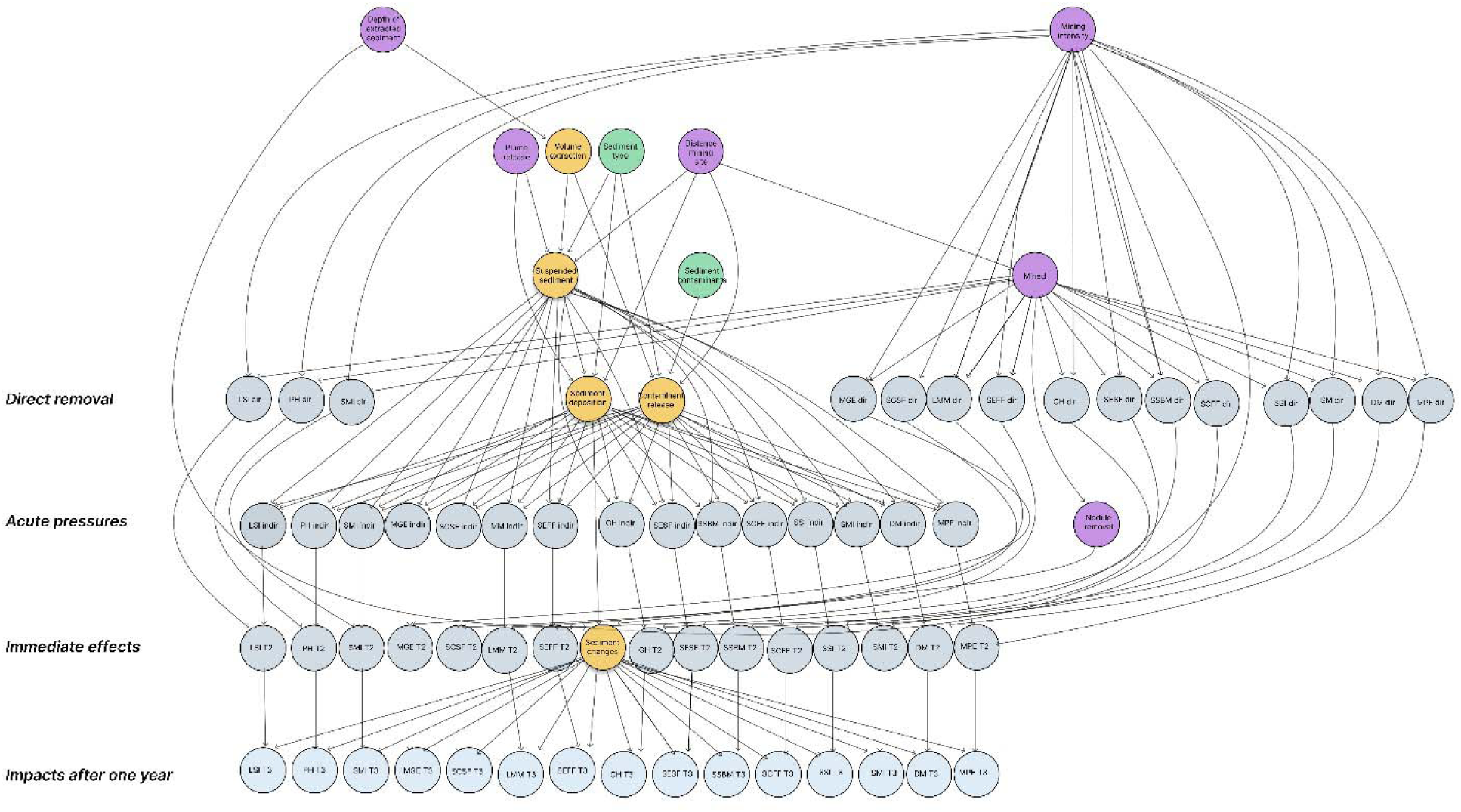
Bayesian Network model for risks of seabed mining on benthic fauna, showing the separate variables for the time steps and all functional groups. Purple circles denote operational variables from the two mining scenarios, yellow circles are pressures arising from mining, green circles environmental conditions (independent of the mining operation), and light blue circles the abundance of the benthic fauna in the different functional groups across the four time-steps in the BN model. For abbreviations of the functional group names, see Table 2.

### 3.1 Likelihood of pressures from mining

The probability of different levels of sediment deposition varied as a function of the distance from the mined area and between the two disturbance scenarios. The most likely outcome under both scenarios was high deposition inside the mining block and low deposition in the far-field (Fig. 5). Under the High disturbance scenario (100% mining intensity inside the mined area; Figure 2, Table 3), the probability of high sediment deposition was estimated to be 0.80 inside the mined area. In the near-field the most likely outcome was moderate sediment deposition with a 0.57 probability, and in the far-field the most likely outcome was low sediment deposition (0.87 probability). For suspended sediment, moderate suspended sediment concentrations (SSC) were the most likely outcome inside the mined area and in the near-field area. In the far-field, low SSC levels were the most likely (0.71 probability). The probability of high SSC was 0.08 inside the mined area, 0.07 in the near-field, and 0 in the far-field.

**Figure 5.**
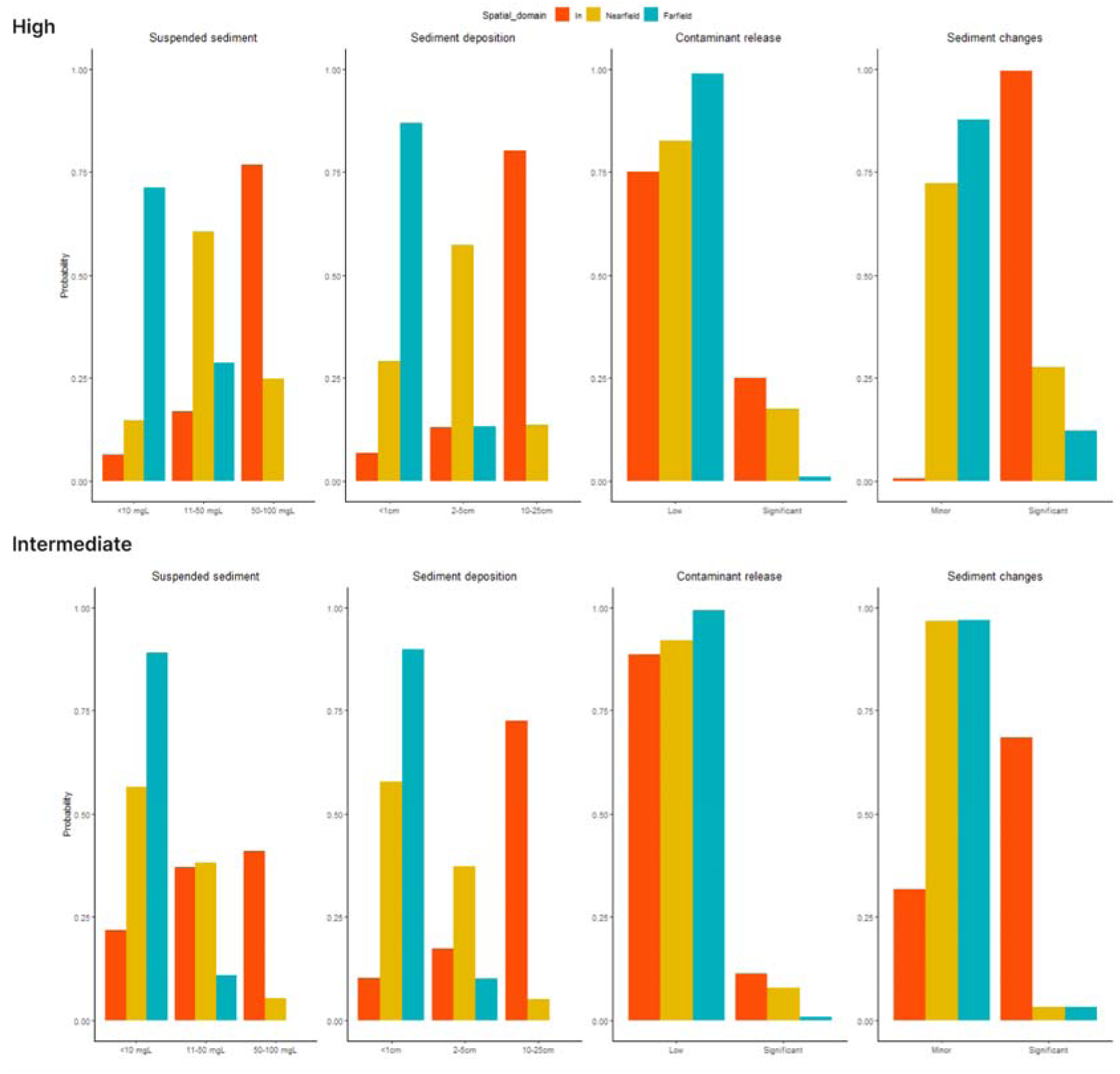
Probability of the different levels of suspended sediment concentration, sediment deposition, contaminant release, and sediment changes resulting from mining inside the mined area, and the near-field and the far-field outside of the mining block under the two disturbance scenarios (High and Intermediate).

The magnitudes of SSCs and sediment deposition were estimated to be lower under the Intermediate disturbance scenario (50% mining intensity to the mined area; Figure 2, Table 3), in which the depth of extracted sediment would be lower than in the High disturbance scenario (Table 3). Under Intermediate disturbance, the near-field area was expected to receive only a low sediment deposition (0.57 probability). Similarly, low levels of suspended sediment were the likeliest outside the mined area, with 0.56 probability of low SSC in the near-field and 0.89 outside in the far-field area. Inside the mining block high levels of SSC and sediment deposition were the likeliest outcomes, with 0.41 and 0.72 probabilities, respectively.

The largest difference between the two evaluated scenarios was the probability of significant sediment changes. Under the High disturbance scenario, significant sediment changes were expected not only inside the mined area (0.99 probability), but also in the near-field (0.23 probability) and in the far-field (0.19 probability). Under the Intermediate disturbance scenario, the probability of significant changes outside the mining block in the far-field was only 0.03.

As the release of harmful substances was set to be mostly driven by potential concentration of toxic substances in sediment and the sediment substrate type in the model, there was only a small difference between the two scenarios for toxin release. The probability of significant contaminant release from the sediment was low under both scenarios, with a 0.25 probability inside the mined area and 0.01–0.17 in the near-field and in the far-field in the High disturbance scenario, and 0.11 probability inside and zero to 0.08 probability in the near- and far-field areas under the Intermediate disturbance scenario.

### 3.2 Impacts on benthic fauna

The various functional groups were assigned differential responses to the direct impacts to, and subsequent recovery from mining, based on data and expert assessments (Table 4). Mobile epifauna and hyperbenthic species are to an extent able to escape the physical disturbance and thus experience lower decreases in abundance from the direct impacts of mining, whereas sessile fauna inside the mining area will be removed by the sediment extraction. Aside from hyperbenthos, meiofauna were estimated to be most tolerant to indirect impacts of mining. Small infaunal species were estimated to experience moderate to high decreases in abundance from indirect impacts even under intermediate disturbance but had moderate recovery potential after one year. Sessile epifauna, such as stony corals, were estimated to experience high decreases in abundance from direct disturbance, and moderate decreases in abundance from indirect disturbance, but recovery will be limited. However, small soft-bodied sessile taxa may have the potential to recolonise the area within one year. In the following section we present a selection of results of the impacts on benthic fauna, conditional on the mining scenarios and the probability of the magnitude of the ecosystem pressures as presented above. Full results on the impacts on all functional groups are contained in the Appendix 2.

#### 3.2.1 Immediate impacts under the High and Intermediate disturbance scenarios

The decrease in abundance immediately after the mining disturbance varied between the functional groups (Figs. 6–7) and with proximity to the mined area. Under the High disturbance scenario, all sessile and infaunal organisms were estimated to experience high relative decrease in abundance inside the mined area (81–100% compared to the pre-disturbance community) (Fig. 6).

**Figure 6.**
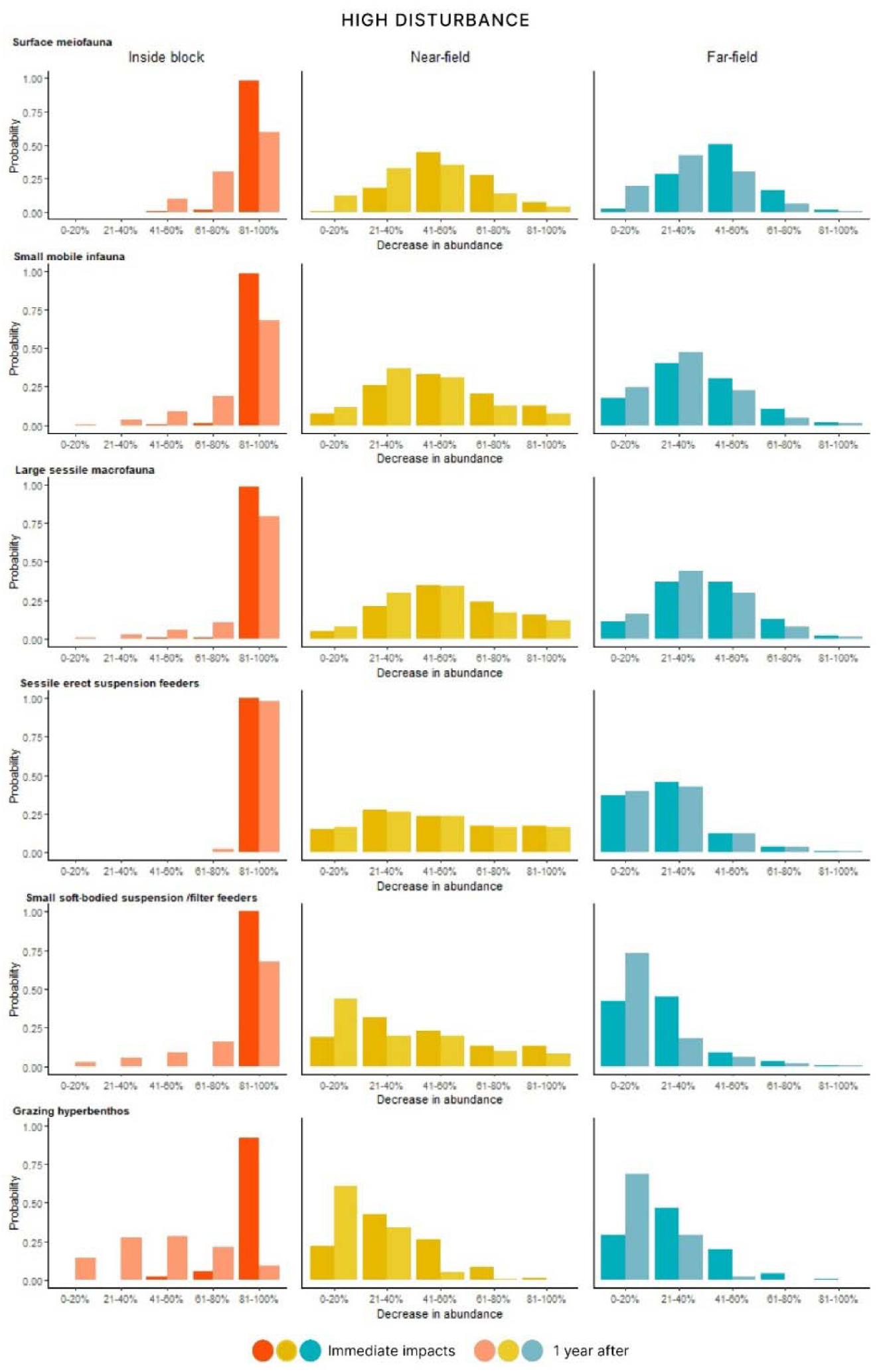
Probability of relative decrease in abundance of six selected epifaunal functional groups inside the mining block (left panel), in the near-field area directly adjacent to the mined area (middle), and in the far-field area outside the mining block (right panel) under the High disturbance scenario. Immediate impacts are noted in a dark shade and impacts after one year in a lighter shade. Full results for all functional groups are contained in Appendix 2.

**Figure 7.**
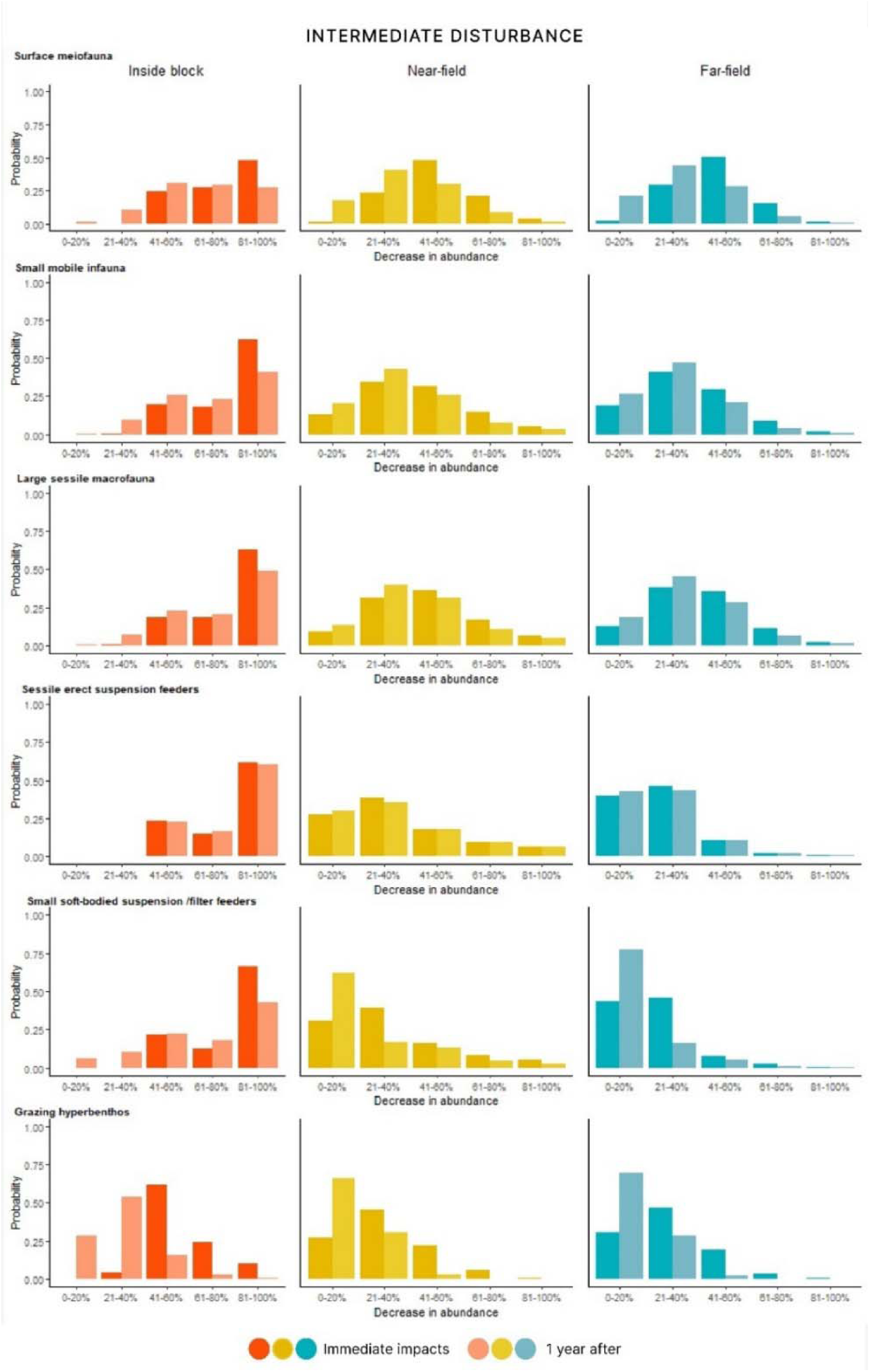
Probability of relative decrease in abundance of six selected epifaunal functional groups inside the mining block (left panel), in the near-field area directly adjacent to the mined area (middle), and the far-field area outside the mining block (right panel) under the Intermediate disturbance scenario. Immediate impacts are noted in a dark shade and impacts after one year in a lighter shade. Full results of all functional groups are contained in Appendix 2.

In the near-field, the most likely outcome under the High disturbance scenario was 41–60% reduction in abundance for all infaunal groups. In the far-field, for most infaunal groups the most likely outcome was 21–40% decrease in abundance. Meiofauna were expected to experience a 41–60% relative decrease in abundance with a 0.5 probability for both deep and surface meiofauna.

Sessile epifauna were estimated to decrease in abundance by 0–60% in the near-field and 0–40% in the far-field, with varying probabilities depending on the functional group. Sessile soft-bodied organisms were estimated to show 21–40% decrease in abundance with a 0.32 probability in the near-field under the High disturbance scenario. In the far-field the most likely outcomes were 20–40% decrease in abundance (0.45 probability), and 0–20% decrease (0.42 probability). Sessile, encrusting organisms were estimated to have higher relative changes in abundance in the far-field and in the near-field than erect forms for both suspension and filter feeders. For example, for sessile encrusting suspension feeders, the most likely outcome in the near-field was 41-60% decrease in abundance with 0.29 probability, whereas for erect suspension feeders the likeliest outcome was 21-40% decrease with a 0.28 probability (full results for all groups in Appendix 2 Figs. S1–S6).

Mobile epibenthos and hyperbenthic organisms had the lowest decrease in abundance in both the far- and near-field areas under both disturbance scenarios (Figs. 6–7), with the smallest decrease in abundance predicted for mobile organisms in the far-field area. For example, for grazing hyperbenthos, the most likely outcome in the near-field and far-field under the High disturbance scenario was 21–40% decrease in abundance with 0.42 and 0.47 probabilities, respectively. Under the Intermediate disturbance, the most likely outcome for hyperbenthos was also 21–40% decrease in abundance with a probability of 0.45 in the near-field and 0.47 in the far-field. Similar to the High disturbance scenario, an 81–100% decrease in abundance was the most likely outcome inside the mined area for infauna and sessile organisms under the Intermediate disturbance scenario. However, a large variation in the impacts inside the mined area was observed under this scenario, with impact estimates for infauna ranging from 40–100% (Fig. 7). In the near-field, for most groups of infauna the most likely outcome under the Intermediate disturbance scenario was 21–40% decrease in abundance. For sessile epifauna, the most likely outcome for all groups was 21–40% decrease in abundance in the near-field and between 0–40% in the far-field with over 0.4 probabilities for each group. Hyperbenthos were estimated to experience a 21–40% decline in abundance in both the near-field and the far-field areas.

#### 3.2.2 Impacts after one year

In our model, the probability of decrease in abundance one year following seabed disturbance is conditional on the initial disturbance-related decrease in abundance and changes in the environment. The estimates of relative decrease in abundance one year after disturbance therefore incorporate both a metric of recovery and any additional decrease due to sub-lethal effects (see Table 4). Recovery was estimated to be more likely across all faunal groups under the Intermediate disturbance scenario, which resulted in overall lower changes in abundance and a lower likelihood of significant sediment changes (Fig. 7). Recovery was more likely to occur in the near-field and the far-field, compared to inside the mined area, as there was less likelihood of significant sediment changes. Similarly, recovery was more likely under the Intermediate disturbance scenario when a larger proportion of the original abundance remained and the sediment changes were smaller, compared to the High disturbance example.

Sessile epifauna had the greatest probability of high relative decrease in abundance (immediate impacts) and were the least likely to show recovery after one year. Under the High disturbance scenario, the probability that 80–100% of their original abundance would still be absent after one year was 0.92–0.97 in the mined area, 0.65 in the near-field, and 0.53 in the far-field for most sessile organisms (Fig. 6). Under the Intermediate disturbance scenario, the most likely outcome inside the mined area for most sessile megafauna was also 80–100% decrease in abundance (Fig. 7), but with smaller probabilities than under the High disturbance scenario (0.50–0.68) (Fig. 6). One exception within this group were the soft-bodied megafauna, which were deemed to experience a 80–100% decrease in abundance inside the mined area after one year with a 0.67 probability under the High disturbance scenario (Fig. 6) and 0.47 under the Intermediate disturbance scenario (Fig. 7). In the near-field, this group was deemed to recover nearly fully (0–20% decrease in abundance) with a 0.43 probability under the Intermediate scenario, whereas under the High disturbance scenario the corresponding probability was 0.26. In the far-field there was 0.56–0.60 probability that there would be no observable change in the abundance of soft-bodied megafauna under either disturbance scenario.

Hyperbenthos were estimated to experience a 20–60% abundance change with a 0.77 cumulative probability inside the mined area after one year. In the near-field the change was estimated to be smaller, with a 0.95 joint probability for 0–40% decrease in abundance. In the far-field the most likely outcome was small or no decrease in abundance (0–20%) with a 0.98 probability. Under Intermediate disturbance scenario changes were much smaller: inside the mined area the most likely outcome for grazing hyperbenthos was 21–40% decrease in abundance with a 0.54 probability. In the near-field and far-field small to no or little decrease in abundances (0–20% change) of hyperbenthos were expected after one year with 0.66 and 0.69 probabilities, respectively.

For small sessile infauna, the probability of 80–100% of the original abundance being lost inside the mining block was 0.74 under the High disturbance (Fig. 6), compared to 0.47 under the Intermediate disturbance scenarios (Fig. 7). In the near-field and far-field, the most likely outcome under both scenarios was a 21–40% decrease in abundance (0.32–0.45 probability under High and 0.40–0.45 under Intermediate disturbance), with a wider distribution with an increasing distance from the mined area and probability distributions converging towards the higher decreases in abundance under the High disturbance scenario. Large mobile macrofauna showed a similar pattern to small infauna, yet with higher recovery rates: in the far-field the most likely outcome was 20–40% negative decrease in abundance with a 0.52 probability under both scenarios.

#### 3.2.3 Comparison of scenarios

Inside the mined area, differences between the two disturbance scenarios were particularly evident for mobile benthos (Fig. 8). In the far-field the differences between the two scenarios were very small for most functional groups, particularly the sessile megafauna. As the probabilities of high impact in the far-field were lower under the Intermediate disturbance scenario, there is less difference between these two spatial domains compared to the differences observed under the High disturbance scenario.

**Figure 8.**
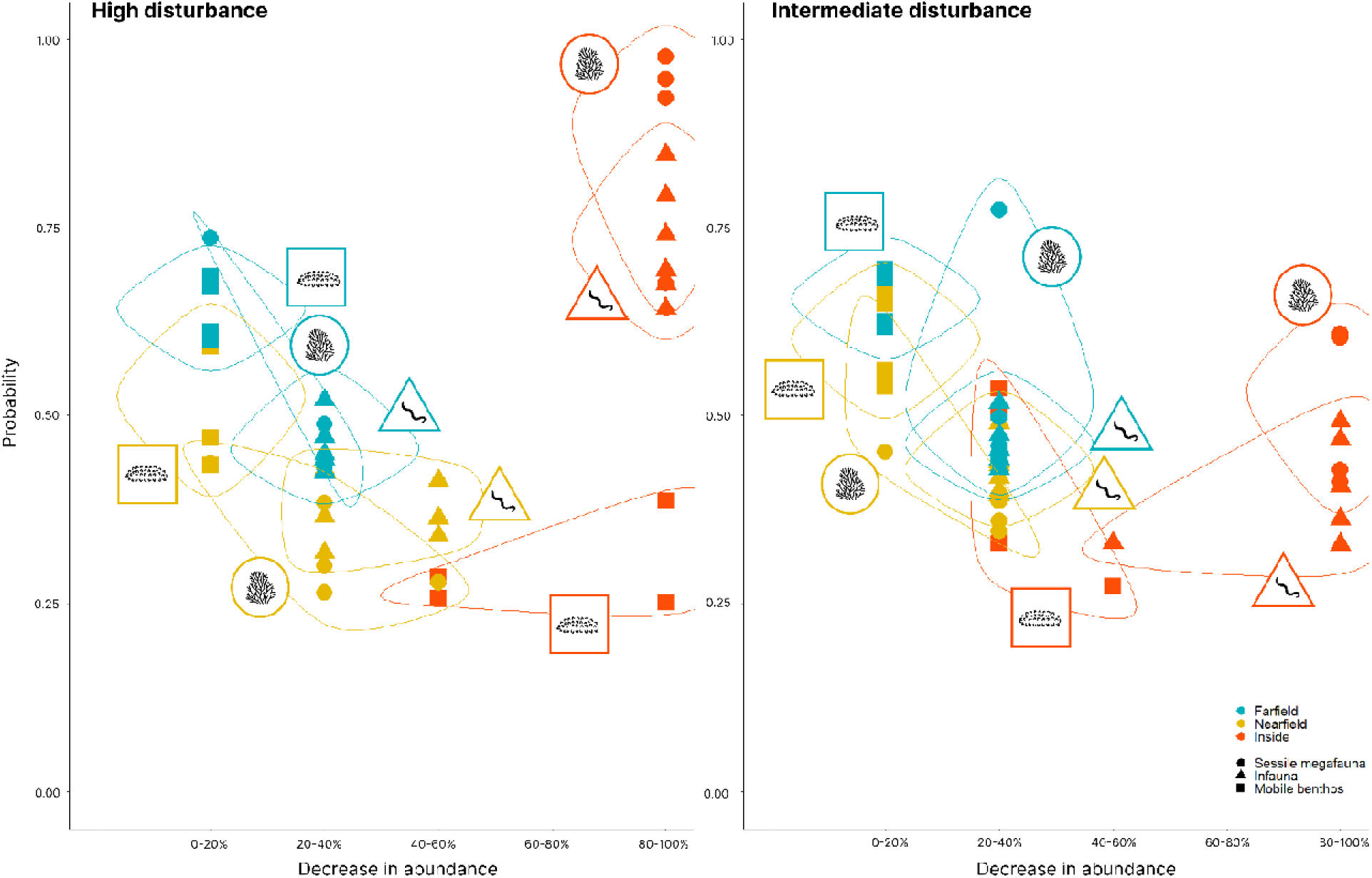
Summary of the most likely outcome for the functional groups in each spatial domain after one year under the High (left panel) and Intermediate disturbance (right panel) scenarios.

Under the High disturbance scenario, the probability estimates for the near- and far-field were more variable. Overall, for both scenarios highest certainties were given to impacts inside the mined area.

The points in the scatterplot represent the most probable outcome for each functional group as a function of its associated probability. The colours depict the three spatial domains (inside minds area, near-field, far-field), and the broad faunal groups (Functional group: sessile megafauna, infauna and mobile) are shown in different shapes and icons.

### 3.3 Model validation and sensitivity

The expert group were satisfied with the BN model’s ability to capture the variation in the impacts across the different seabed mining disturbance scenarios and spatial domains. Reviewing the model results showed that mining intensity had the highest impact on the magnitude of physicochemical pressures. As both the model structure and the parameters are largely expert informed, there was no need to perform a numerical sensitivity analysis to evaluate which variables had the highest impact on the final outcome (as reflected in a decrease in abundance of benthic fauna).

The use of probability distributions in BNs allow the uncertainty regarding the variation in a variable of interest to be directly embedded in the impact estimates. To account for the uncertainty resulting from the lack of information on the process being evaluated we also recorded the experts’ certainty in the probability estimates (Table 5).

**Table 5.** Overview of the confidence in the variable parameterisation. Full details of information used is included in Appendix 1.

| Variable group | *In situ* data | Experimental data | Literature | Confidence |
| --- | --- | --- | --- | --- |
| Sedimentation (SSC and sediment deposition) | Yes | No | Yes (Lescinski et al., 2014) | Moderate |
| Contaminant release | No | No | Yes (Frontin-Rollet 2017; Hauton et al., 2017; Simon et al., 2022) | Low |
| Mobile megabenthos and hyperbenthos | No | No | None (baseline information in Lörz, 2011). | Low |
| Meiofauna | Yes | No | None | Moderate |
| Macroinfauna | Yes | No | None | Moderate |
| Sessile megafauna | No | Yes | Yes (Bell et al., 2015; Brooke, Holmes, and Young 2009; Leys 2013; Mobilia et al., 2021; 2023; Pineda et al., 2017; Wurz et al., 2021) | Moderate |

For the physicochemical parameters, highest uncertainties were assigned to potential release of harmful substances from the sediment and extracted phosphatic mineral material (Table 5). Similarly, the impacts of toxin release on all groups of benthic fauna were ranked as highly uncertain, as few studies have been published on the topic and the number of potentially harmful substances is unknown. For this reason, the importance of toxin release received a low weight in the impact estimates for all benthic functional groups, but this should be considered in future ERAs as a potentially significant stressor.

The highest uncertainties within functional groups were assigned to mobile epibenthic organisms and hyperbenthos (Table 5). This uncertainty is also reflected in the probability estimates, where hyperbenthos estimates are the most varied (broadest distribution). In turn, meiofauna and small mobile infauna received highest certainty, as for these taxa both new data and previous studies could be used to assess the impacts (Tables 2 and 5).

## 4 Discussion

Uncertainty regarding biological responses of marine organisms to seabed disturbance is a major concern for estimating the impacts of human activities in the deep sea, which subsequently impacts upon decision- and policy-making (Kung et al., 2021). To quantify the uncertainties in biological responses, we developed a probabilistic ecological risk assessment model to describe the pressures caused by deep seabed mining and the responses of affected benthic ecosystem components. We estimated the magnitude of the ecosystem responses and probability of recovery by combining field and experimental data, information from published literature, and expert knowledge. This type of model can be used to identify the likelihood of ecosystem losses from seabed disturbance to guide the regulation and management of such activities.

The BN model successfully captured the variation in the likelihood of disturbance from the two different disturbance scenarios assessed. As the highest pressures were confined to inside the mined area, largest differences between the scenarios were seen inside the mined area and in the near-field adjacent to the mining block. Mining intensity and depth of sediment extraction were the key drivers of the pressures arising from mining, and there was a logical spatial gradient within the two mining scenarios. In the High disturbance scenario, the model predicted high levels of suspended sediment and sediment deposition to be unlikely outside the immediate area of disturbance. Despite the moderate to low levels of physicochemical disturbance, most benthic organisms, regardless of their functional group, were predicted to decrease in abundance by 60– 100% inside the mined area and by 20–60% in the unmined near-field. The highest levels of relative changes in abundance in the far-field were for encrusting sessile suspension and filter feeders. Under the Intermediate disturbance scenario, changes in abundance were lower and there was more variation in the potential responses of fauna inside the mined area. In the near-field under this scenario, 20–40% decrease in abundance was the most likely outcome for most groups. In the far-field area the differences between the two scenarios were smaller; overall, after one year, mobile organisms were expected to experience a 0–20% loss in abundance, whereas for all other organisms 21–40% decreases were expected. Importantly, we noted increasing levels of uncertainty in the estimates for biological responses with increasing distance from the mined area and relatively lower levels of pressures from mining.

Most functional groups evaluated in this study were anticipated to tolerate low levels of sedimentation relatively well. As high levels of sediment deposition were estimated to be confined to inside the mining area, most organisms were predicted to experience a decrease of less than 60% of the original community in the near-field and far-field under both scenarios. However, it is important to note that the estimates here reflect the characteristics of seafloor communities in our case study area, and are not necessarily directly applicable to other areas, such as in regions where abyssal manganese nodules (e.g., Clarion-Clipperton Fracture Zone) or placer or dredged deposits may be extracted from the seafloor (e.g., offshore Namibia).

The use of separate time steps allowed us to quantify how different organisms react to changes in their environment directly after mining and one year later, incorporating a simplified metric of recovery and sub-lethal effects, which are potentially important for the mortality of larger organisms (e.g., Martins et al., 2022). While some groups of organisms had similar responses to the immediate effects inside the mined area (i.e., small infauna and sessile organisms), the impact estimates after one year showed that mobile fauna and small-sized fauna experienced fewer negative impacts than large infauna and long-lived sessile megafauna. Sessile organisms showed little to no recovery within this one-year timeframe (also found in the review by Jones et al., 2017).

A key underpinning factor leading to the rejection of the CRP mining consent application was that it did not quantify the scale of effects on benthic communities away from the mining blocks (NZ EPA 2015). In this model, the inclusion of discrete spatial domains allows the impact of disturbance on the benthic community to be assessed quantitatively under different disturbance regimes inside the mined area, in the near-field (adjacent to the mining block), and in the far-field (further away but still within the impact zone). However, the assumption was made that all mining would be completed within the mining block. The detailed pattern of mining (e.g., blocks, strips) was not factored into the analysis and we assumed homogeneous impacts inside each spatial domain. An important addition to this approach that would make it more directly relevant to a proposed operation would be combining this model with spatially explicit data, such as sediment plume modelling, to estimate the spatial extent of the suspended sediment concentrations and sediment deposition from the mining activities as a function of distance from the mined area (Lescinski et al., 2014; Spearman et al., 2020). A similar approach can be used if detailed spatial information on the benthic community is available (e.g., Helle et al., 2020), enabling a more precise spatial representation of potential impacts. It is important to note that high uncertainties remain regarding effects on mobile taxa. An improved understanding of impacts on epibenthos, hyperbenthos, and fishes in future research is essential to fully assess the extent and magnitude of mining or sediment disturbance (Washburn et al., 2023).

### Incorporating ecological data into ERAs

Despite an increase in research regarding the impacts of seabed disturbance to seafloor ecosystems (e.g., Gollner et al., 2017; Jones et al., 2017), some experts consider there are few categories of scientific knowledge comprehensive enough for all the relevant ecosystem components to enable evidence-based decision-making and robust environmental management of such activities (Amon et al., 2022). To overcome the inherent data paucity in many deep-sea environments, it is necessary to use all possible scientific evidence, from other industries (e.g., Kaikkonen et al., 2018) as well as analogies to shallow-water systems and communities (Van Der Grient and Drazen 2021). Our approach in combining empirical data and expert assessment demonstrates that, despite the considerable body of literature on the different aspects of physical and sedimentation impacts, formulating conclusions on the impacts is not an easy task. In light of these challenges, the probabilistic approach as employed here proved useful, as the uncertainties related to the impacts were directly incorporated in the impact estimates. We found that experts were more comfortable giving uncertain judgements when this aspect was embedded in the process. Furthermore, the approach provides a method to synthesise information from multiple sources and move from qualitative risk statements to more quantitative impact estimates. As the conditional probabilities may be drawn from multiple sources, the model can be continuously updated as new information becomes available (see Table 5 for used information sources and evidence gaps).

A major issue with determining the sensitivity of species or groupings of functionally similar organisms to environmental disturbance is that most deep-sea organisms are poorly studied, thus detailed information on the ecological responses is limited outside some specific habitats such as hydrothermal vents (Chapman et al., 2019). Nevertheless, trait-based approaches are useful in identifying species responses to direct anthropogenic impacts and other environmental changes (Baird, Rubach, and Brinkt 2008; Boschen-Rose et al., 2021; Krumhansl et al., 2016; Lundquist et al., 2018). We found that generalising biological responses using broad functional groups allows for a pragmatic understanding of how organisms may respond to external stressors, which could improve development of options for effective management and conservation strategies (Miatta, Bates, and Snelgrove 2021). The use of broad functional groups facilitates application of data collected from one area to another, although in such cases, the variation in the trait expressions associated with biological responses across regions must be carefully evaluated (de Juan et al., 2022).

### Improving probabilistic risk assessments

Human activities may result in multiple complicated changes in the environment with different spatial and temporal scales, and as a result, modelling such complex systems with any method comes with drawbacks and often requires simplification to enable a satisfactory result (e.g., Uusitalo 2007). Handling a large number of variable connections, impact pathways, and associated evidence is a demanding task, and ensuring that all experts participating in the assessment understand the study background is a challenge. The increasing complexity of integrating different types of data and knowledge (biological, technical, geological, geotechnical, oceanographic) into the chosen model not only poses communication challenges for the assessment team and between experts, but handling and organising the collected information requires considerable effort by the experts and the modellers. Model complexity also adds to the workload for parameterisation of the model, and using an elicitation tool for filling the CPTs is often the only feasible solution when there are many variables to quantify (Pollino et al., 2007). However, the use of a parameter-based estimation method (such as the beta interpolation used in this study), while easy to carry out, may produce estimates that do not fully reflect the expert’s complex views on a topic.

To avoid producing incorrect estimates in this BN modelling, all CPTs generated with the interpolation tool were reviewed both by the expert team and the modeller coordinating the effort. When these caveats are considered in the quantification process, the use of interpolation tools provides a useful option for evaluating the effects of multiple stressors, which would otherwise be impossible to quantify due to the high number of possible probability entries required. Therefore, while the workload involved in quantifying a large BN model may seem overwhelming, a similar (or larger) workload applies for any kind of impact assessment where the complex connections between components are being assessed. Another drawback of using BNs is their acyclic nature, preventing the inclusion of feedback loops between parent and child nodes (e.g., for ecosystem interactions; Uusitalo, 2007). This issue can be partially overcome with the use of splitting nodes, or through Dynamic BN approaches with multiple time steps (Trifonova et al., 2015).

There are several ways the BN model developed here can be augmented to better estimate the potential impacts from seafloor disturbance. First, improving recovery estimates of the geochemical and biological components in the model would be beneficial to account for the magnitude of the disturbance in the recovery estimates. Even if the pelagic realm remains more poorly studied than seafloor ecosystems (Bisson et al., 2023; Robison 2004), broadening impact estimates to account for water column impacts and including other groups of organisms, such as micro-organisms (Herndl and Reinthaler 2013) and fishes (Drazen et al., 2020), would allow for more holistic estimation of impacts. Similarly, although our method expands the time horizon of seabed mining impacts from acute to longer-term impacts by including recovery potential over one year, the approach does not include detailed information on the full interactions within the ecosystem. For example, long-term (multi-decadal) deleterious impacts on ecosystem functioning have been demonstrated in disturbance experiments in abyssal nodule fields (e.g., Peru Basin, Vonnahme et al., 2020). As a result, to improve our method, it will be useful to more carefully examine the interplay between different ecosystem components, such as competitive food web connections or biogeochemical linkages. The approach further allows the integration of cumulative impacts in the risk assessment, e.g., to account for climate change effects (Furlan et al., 2020). Finally, to overcome the simplification required when using discrete variables, moving to hybrid networks that allow mixing continuous and discrete variables would provide opportunities to describe the impacts more precisely (e.g., Moe et al., 2020).

### Further applications

Environmental management often requires decision-making under uncertainty regarding the potential outcomes of activities and the most effective ways to mitigate them. Despite recommendations for the use of probabilistic methods in risk assessments (Van den Brink et al., 2016), their comprehensive integration into regulatory risk frameworks is still limited. Deterministic approaches, such as calculating single risk values based on a predicted exposure to a stressor remain more prevalent (Fairbrother et al., 2016). By utilising a probabilistic model capable of generating estimates for various scenarios, it would be feasible to identify management actions that are most likely to minimise stressor inputs in the case of deep-sea mining, leading to improved chances for the maintenance of the ecological functions of impacted deep-sea faunal communities.

The probabilistic model described in this study was developed, from a scientific perspective, to provide a framework for further applications. For a real-world application for management purposes, it is important to engage with the relevant regulatory bodies and stakeholders to ensure that the model framing and metrics align with societal and management needs (e.g., specific species and habitats or maximum thresholds for allowed impacts). For management purposes, a useful quality of BN models is that they may be further augmented to incorporate socioeconomic data (Uusitalo et al., 2022). To ensure the optimal use of the models, such risk assessments should involve interdisciplinary collaboration between a diverse group of scientists, policymakers, and stakeholders to ensure that the best available knowledge is integrated into the decision-making process.

There are important considerations when applying the approach and the results presented here to other deep-sea ecosystems and forms of disturbance. While the results give some insights to the broad patterns of how different functional groups of deep-sea organisms may respond to seabed disturbance, the magnitude of pressures and the responses of biological communities is likely to vary considerably from one area to another depending on the prevailing environmental conditions, connectivity of seafloor communities, and the types of disturbance (e.g., Boschen et al., 2013; Haffert et al., 2020; Jones et al., 2017). Based on the combined laboratory experiments (Mobilia et al., 2021; 2023) and field data, the biological communities on the Chatham Rise may be considered to be better adapted to temporary increases in suspended sediment that might typically be experienced under the Intermediate disturbance scenario than those in more stable systems (such as abyssal plains). However, the results of the model indicated that most benthic functional groups of the Chatham Rise were expected to decrease in abundance and were not expected to recover even from Intermediate disturbance outside the mined area after one year. The results suggest high uncertainties regarding the impacts, especially outside mined seafloor areas, and stress the importance of further studies on the recovery dynamics at broader spatial and temporal scales. Applying similar quantitative risk assessment models in other areas where deep-sea mining is considered, such as the Clarion-Clipperton Zone, is therefore important, as it enables a systematic and data-driven evaluation of potential risks, environmental impacts, and uncertainties across multiple habitats associated with this emerging industry.

## Acknowledgements

We thank the ship and science crews of the NIWA voyages TAN1805, TAN1903, and TAN2005 and the NIWA staff involved in processing the field data. Di Tracey, Chris Hickey, Savannah Goode, and Katie Bigham are acknowledged for their valuable insights on the model parameterisation. We also thank Laura Uusitalo, whose advice in Bayesian Network modelling greatly improved our analysis. Funding for this work was provided by Ella and Georg Ehrnrooth Foundation, Maj and Tor Nessling foundation, and New Zealand Ministry of Business, Innovation and Employment (MBIE contract CO1X1614).

