## Appendix 1 for "Probabilistic ecological risk assessment for deep-sea mining: a Bayesian Network for Chatham Rise, SW Pacific Ocean"

- Appendix 1 S1 Detailed method description
- Appendix 1 Table S1 Example of a conditional probability table
- Appendix 1 Table S2. Experts involved in the modelling process & probability elicitation
- Appendix 1 Table S3. Details of variable discretisation and parameterisation

### S1 Methods description

#### Step 1: Participatory modelling

The first stage of the work focuses on defining which ecosystem components should be included in the risk model and mapping out their interdependencies based on a previous network drawn from literature. This consisted of defining the model structure with experts in semi-structured interviews, which were implemented as a causal mapping exercise, asking experts to identify the impact pathways associated with seabed mining. We held both joint workshops and individual interviews, which were complement by data analysis from the field measurements as well as literature reviews. Model building consisted of first identifying variable related to the mining technology to recognise the pressures caused by mineral extraction, then the relevant environmental conditions that may affect the magnitudes of these pressures, and finally the affected non-biological (e.g., changes in seabed topography, sediment composition), and biological ecosystem components. This step resulted in a generic modelling framework for any type of benthic organism.

#### Step 2: Variable selection: Assigning functional groups

Based on the general causal network developed in step 1, we defined variables describing the benthic ecosystem components that were likely to be affected by the disturbance from mining. As the number of individual species on the Chatham Rise is far too high to assess each species or taxa separately, we reduced this complexity by grouping organisms into functional groups. The functional groups were assigned based on the expected response of organisms to the different pressures caused by mining, using previously created groupings for the Chatham Rise as a starting point (Lundquist et al., 2018). Functional groups in the model were selected based on traits that have been shown to affect impacts to sedimentation and recovery potential of organisms: body size, feeding habit, position in sediment, and mobility (Hewitt, Lundquist, and Ellis 2018). The number of trait expressions used to determine the groupings varied between functional groups. If a trait was not seen to affect the response in some broader group of organisms, all the different combinations of trait expressions were not used.

The final causal network was modelled as a Bayesian network (BN). Bayesian statistics provide an alternative to commonly used simple scoring procedures in ecological risk assessment. For an introduction to BNs, see e.g., Kaikkonen et al. (2021), Aguilera et al. (2011) and Chen and Pollino (2012).

**Step 3: Variable discretization**

Discrete variables were defined to describe the variation in the magnitude of pressures arising from the mining activity. This included considering which combinations of pressures the extraction and sediment deposition are likely to result in. The discrete states may be described through quantitative metrics, like different concentrations of substances or depth in centimetres, or can be based on qualitative descriptions of discrete classes (e.g., high, medium, low). The key aspect with regard to the biological responses was that these pressure levels should make sense from an ecological perspective. As the demonstration in our case study is not bound to a specific setting, we decided to frame the model variables quite generally with the states low-moderate-high. The discrete variable states were mostly defined based on expert judgement, informed by data and literature (see Table S2 for details).

**Step 4: Model parameterisation**

Within a BN, the magnitudes of impacts are illustrated through conditional dependencies. The probabilities of each value of the child node, conditioned on every possible combination of values of the parent nodes, were drawn from expert opinion. These describe the strength of the causal relationships between variables in the model.

All conditional probability tables (CPT) were developed manually by experts using field and experimental data and literature as supporting information (see Table S2). The following process was followed for the different variables.

We used the graphical interface provided in the Application for Conditional probability Elicitation (ACE) application (Hassall et al., 2019) to initialise the CPTs for the geological variables. The application provides a starting point for defining the overall shape of a conditional probability distribution by ranking the direction and magnitude of the parent nodes on the child node and populating the table through a scoring algorithm. For the probabilities concerning the impacts of direct pressures on benthic fauna, the prefilled tables were evaluated and adjusted in another session with all experts to reach a consensus on the magnitude of the impacts.

For variables describing the impacts on benthic fauna, direct and indirect mortality were modelled separately. This division accounts for the total mortality of benthic fauna, which encompasses both direct mortality resulting from sediment and mineral concretions extraction, and indirect mortality arising from other pressures such as sediment deposition, suspended sediment, and the release of harmful toxic substances. This approach enables us to estimate the effects of these pressures in both the direct mining area (total mortality) and neighbouring areas (indirect mortality). Organisms within the mining block will experience both direct mortality and, in cases where <100% of the block is mined, the remaining fauna inside the block will be affected by sedimentation. Organisms in the near-field and the far-field will only be impacted by sedimentation effects. This separation was achieved by introducing three auxiliary variables to describe changes in faunal abundance:

- Direct impacts
- Indirect impacts
- Total decrease in abundance

The model was set so that direct impacts applied only inside the mining block, and indirect impacts in all domains. We calculated the probability of changes in abundance from the indirect impacts given the initial direct removal inside the mining block using equation (1). Outside the mining block (in the near-field and far-field) all changes in abundance are due to the indirect impacts.

Direct mortality was estimated as a direct proportion of the mined area, so that e.g., mining 50% of an area results in 50% of fauna being extracted.

For the indirect pressures, we elicited the impacts on each faunal group using a combination of data when available and expert knowledge (Table S2). In order to reduce the elicitation burden on experts, we applied an interpolation method (Barons et al. 2022) to derive the probability distributions. The method assumes that the child node can be estimated through a beta distribution and requires the user to identify both a "best case" and a "worst case" distribution for the child node. In this context, "best" and "worst" pertain to distributions where the parents affecting the node are at their highest and lowest points for the child, respectively. The probability distributions for all other combinations of parent variables are then inferred by interpolating between these extremes. This procedure is accomplished using a series of weights assigned to the parent states, which are employed to interpolate between the parameters of a beta distribution.

We elicited the probabilities for the best- and worst-case scenarios using the following questions:

- What is the lowest the value could be?
- What is the highest the value could be?
- What is the most likely value?

For the biological variables describing the impacts on functional groups for which no data were available, we referred to existing data sets from the Chatham Rise to estimate the approximate community composition of the benthic community within the selected functional groups (e.g., the proportion of demosponges-glass sponges) to assist in the probability estimates. Despite the use of the CPT interpolation tool, all CPTs were carefully reviewed to ensure that the experts' views on the potential impacts were correctly captured by the weights given to the parent nodes.

The separate CPTs for direct and indirect impacts were combined so that the total mortality of benthic fauna within a discrete block and one moment in time comprises the direct mortality from extraction of sediment and mineral concretions, and the indirect mortality of the remaining fauna that are exposed to the pressures from the extraction activity (i.e. one organism cannot be both extracted and die of sediment deposition). The probability of total mortality of benthic fauna was thus calculated as done in (Kaikkonen, Helle, et al., 2021):

$$(1)\ P(Total\ mortality) = P(Direct\ Mortality) + P(Indirect\ Mortality)\ x\ (1 - P(Direct\ Mortality))$$

where p(Indirect Mortality) x (1-p(Direct Mortality)) accounts for the probability of the proportion of fauna remaining after direct extraction. While the resulting joint probabilities are continuous, here we calculate them at 1% accuracy (e.g., round them up) to provide the probabilities of a mortality of a given proportion of benthic fauna for the five discrete classes we use in our model.

**Table S1.** Example of a part of a finalised conditional probability table (CPT) for indirect effects on benthic fauna. The CPT summarises all possible combinations of the different variable states for a given node (full table has 18 rows for all the combinations of parent variable states).

| Suspended sediment | Sediment deposition | Contaminant release | Decrease in abundance | | | | |
|---|---|---|---|---|---|---|---|
| | | | 0-20% | 21-40% | 41-60% | 61-80% | 81-100% |
| Low | Low | Insignificant | 0.8928 | 0.0735 | 0.0235 | 0.0095 | 0.0007 |
| Low | Low | Significant | 0.8019 | 0.1297 | 0.0548 | 0.0126 | 0.001 |
| Low | Medium | Insignificant | 0.8306 | 0.1058 | 0.0411 | 0.0156 | 0.0069 |
| Low | Medium | Significant | 0.7001 | 0.1774 | 0.0784 | 0.0354 | 0.0087 |
| Low | High | Insignificant | 0.0146 | 0.1999 | 0.3378 | 0.2561 | 0.1916 |
| Low | High | Significant | 0.0115 | 0.0861 | 0.2207 | 0.3557 | 0.326 |

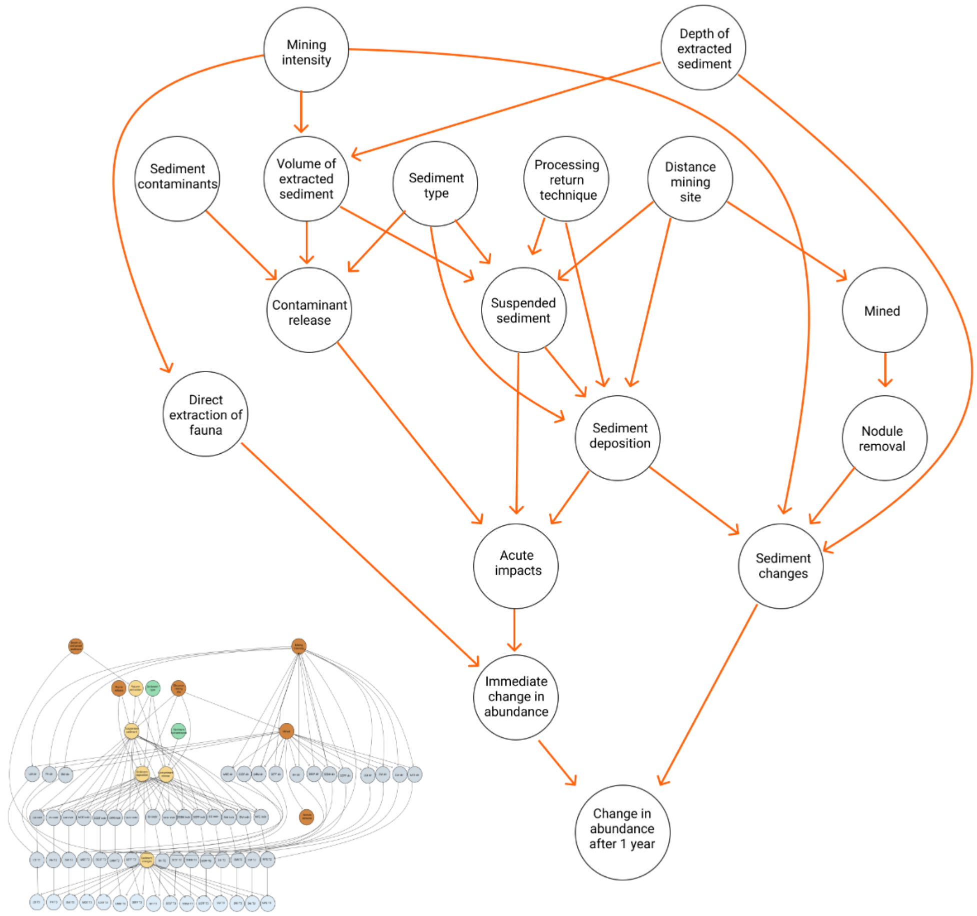


**Figure S1**. Example of a Bayesian network structure for one functional group.

While the uncertainties in the estimates are usually thought to be embedded in the probability estimates (i.e., the wider the distribution, the lower the confidence on the estimate is thought to be), we decided to use an additional metric of certainty, as the probability estimates also encompassed the variability in the functional groups consisting of several species with differential responses to disturbance and recovery.

#### Step 5 Application to scenarios

To evaluate the effect of changes in the mining operations on the impacts on benthos, we defined two alternative mining scenarios. These scenarios, which we define as a combination of specific states of the decision variables that describe the overall mining process are assumed to be controlled by the party responsible for the mining operation. We chose to evaluate two scenarios: one characterising the most likely mining operation on the Chatham Rise, and another more similar to surface collectors that may be used for abyssal deep-sea mining. The scenarios were defined by experts based on the description of the planned mining activities on the Chatham Rise (Chatham Rock Phosphate, 2014). All the other variables in the model are further affected by these decision variables (Fig. S1). It should be emphasised that the model can be queried for any combination of variables, and we have presented only a limited number of possible outcomes. The BN model was queried on these two scenarios by setting the values of the decision variables (details included in the R script available at https://github.com/lkaikkonen/CR-ERA).

**Step 6 Model evaluation**

The model outcomes were evaluated in a series of meetings with the expert group and some of the CPTs were adjusted to better reflect the expected outcomes from the mining disturbance especially regarding the spatial scale of the impacts.

**Table S2**. Experts involved in the expert elicitation for providing probability estimates and/or supporting information for the model parameterisation. The specific fields of expertise are not comprehensive.

| **Name** | **Expertise** | **Model variables contributed to** | **Institute** |
|---|---|---|---|
| Ashley Rowden | Marine Ecology | Infauna | National Institute of Water and Atmospheric Research and Victoria University |
| Daniel Leduc | Marine Ecology | Infauna | National Institute of Water and Atmospheric Research |
| David Bowden | Marine Ecology | Mobile epibenthos and hyperbenthos | National Institute of Water and Atmospheric Research |
| Vonda Commings | Marine Ecology | Sessile megafauna | National Institute of Water and Atmospheric Research |
| Jennifer Beaumont | Marine Ecology | Sessile megafauna | National Institute of Water and Atmospheric Research |
| Di Tracey | Marine Ecology | Sessile megafauna | National Institute of Water and Atmospheric Research |
| Savannah Goode | Marine Ecology | Sessile megafauna | National Institute of Water and Atmospheric Research and Victoria University |
| Scott Nodder | Marine geology and biogeochemistry | Sediment changes, suspended sediment, sediment deposition | National Institute of Water and Atmospheric Research |

**Table S3.** Details of variable discretisation and parameterisation

| Variable name | Description | Variable states | Discretisation method | Parameterisation method |
|---|---|---|---|---|
| Depth of extracted sediment | Depth of sediment extracted by the mining tool | <10cm / 10-30 cm / >30cm | Variable states were defined by experts based on literature to describe different potential seabed mining operations and to be applicable to other types of seabed disturbance, such a bottom trawling. The lowest class describes surface collector operations or low-penetration bottom trawling operations, 10-30cm traditional bottom trawling (Eigaard et al., 2016; Muñoz-Royo et al., 2022), and the >30cm the most likely extraction depth for the planned phosphorite mining (Chatham Rock Phosphate, 2014). | Not applicable |
| Processing return technique | Depth of processing return water and sediment | 10m from seafloor / at the seafloor | Discrete classes were retrieved from the Chatham Rock Phosphate mining consent application (Chatham Rock Phosphate, 2014). | Not applicable |
| Mining intensity | Proportion of area mined (or disturbed in a non-mining application) within a discrete mining block | 50% / 75% / 100% | Three discrete classes were decided by experts to show potential variation in the mining intensity and to align with description given in the Chatham Rock Phosphate Marine Consent Application (Chatham Rock Phosphate, 2014). | Not applicable (decision variable) |
| Distance from the mining block | Distance from the mining block | Inside mining block / Near-field / Far-field | The impacted area was split into three zones based on expert opinion and available results in the Chatham Rock Phosphate Marine Consent Application (Chatham Rock Phosphate, 2014). | Not applicable (decision variable) |
| Volume of extracted sediment | Volume of sediment removed by a mining operation tool (as | Low-medium-high | Volume of sediment was not given a numerical value due to the conceptual nature of the model and lack of exact information on the volumes to be | Literature, expert assessment |

| Variable name | Description | Variable states | Discretisation method | Parameterisation method |
|---|---|---|---|---|
| | the initial removal) | | extracted in literature. The three relative states were chosen by experts using available information in the Chatham Rock Phosphate Marine Consent Application (Chatham Rock Phosphate, 2014). Note that the low-med-high will be relative to the type of seabed operation to be evaluated with the model (mining, trawling etc.) | |
| Particle size composition of the extracted sediment | Proportion of fine and coarser particles of the extracted substrate and in the composite sediment plume | Mostly silt and clay (fine) / mix of fine and coarse particles / mostly coarse (sand and gravel) | Field data from the area | Field survey particle size data before and after disturbance was used to inform probability distribution. |
| Sediment contaminants | Concentration of potentially harmful substances in the sediment and the mineral material to be extracted | Low-Medium-High | Three relative classes decide by experts due to high number of potential biologically harmful contaminants and lack of information on their presence (Frontin-Rollet, 2017; Hauton et al., 2017). | Probability estimates provided by experts using literature as supporting material |
| Nodule removal | Proportion of phosphorite nodules removed from a discrete mining block. | Yes / No | Not applicable | Not applicable (decision variable) |
| Suspended sediment | Total suspended solids concentration near the seafloor resulting both from the processing return | Low: <10 mg $L^{-1}$ / Moderate: 11-50 mg $L^{-1}$ / High:>100 mg $L^{-1}$ | Three discrete classes were chosen to align with the SSC levels used in the lab experiments, levels detected in the disturbance experiment and baseline studies, and modelled SSC values from the sediment plume modelling dispersal for Chatham Rock Phosphate | Conditional probabilities were estimated by experts using the field data measurements and modelled values on the SSC dispersal as supporting |

| Variable name | Description | Variable states | Discretisation method | Parameterisation method |
| --- | --- | --- | --- | --- |
| | and mining tool operation | | (Lescinski et al., 2014) to deliver a contrast in biological responses in terms of extraction impacts. | information. (Lescinski et al., 2014). |
| Sediment deposition | Depth of sediment deposited from the composite suspended sediment plume | Low: <1-2 cm / Moderate: 2-5 cm / High:10-25cm | Natural sedimentation rates estimated from Lander data picking up SCIP disturbance compared with mooring data (for natural sedimentation rates) gave inform this. Likely deposition rates from mining operations and relevant thresholds for benthic fauna from (Chatham Rock Phosphate, 2014; Hewitt & Lohrer, 2013) | Field measurements and expert assessment |
| Contaminant release | Release of contaminants from the sediment plume to the seabed water column | Nonsignificant / Significant release of contaminants | Due to scarce evidence on the release of contaminants and their effect on benthic fauna, a binary variable was deemed most appropriate. | Probability was based on expert assessment, only a marginal probability was given for significant response due to lack of evidence. |
| Changes in sediment characteristics | Measure of changes in the sediment environment affecting habitat quality for benthic organisms, including changes in:<br>• Grain size<br>• Sediment porosity | Minor to no changes / Significant changes | Due to scarce evidence on the release of contaminants and their effect on benthic fauna, a binary variable was deemed most appropriate. | Field measurements data on sediment particle size change was used to calculate relative change in mud content. Total sediment change was estimated as the combination of nodule removal and sediment particle change and final probabilities were given by experts based on the field data. |

| Variable name | Description | Variable states | Discretisation method | Parameterisation method |
|---|---|---|---|---|
| | • Sediment compaction<br>• Organic enrichment | | | |
| All benthic fauna | Relative change in abundance of benthic fauna | | Five discrete classes decided by expert team to divide the continuous abundance change to provide a comparison and to limit the number of required probability entries to facilitate probability elicitation | Field measurements, used to inform the probability estimation. Probability estimation done with experts using and reviewed with the entire expert groups to ensure consistency between estimates for different faunal groups. |
