## Appendix 2 for "Probabilistic ecological risk assessment for deep-sea mining: a Bayesian Network for Chatham Rise, SW Pacific Ocean"

- Appendix 2 Figures S1-S6: Full results of model runs under different scenarios

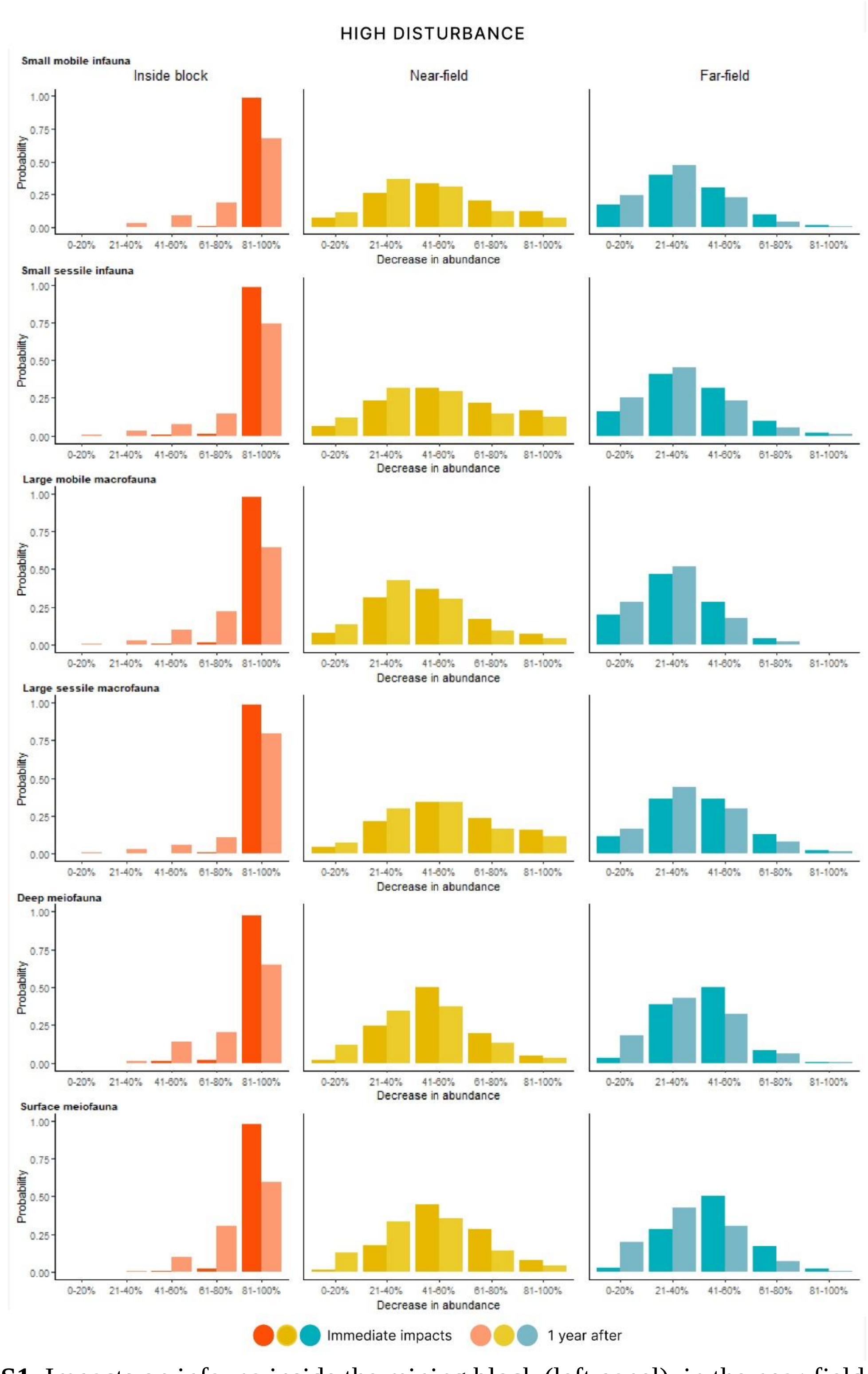


**Figure S1.** Impacts on infauna inside the mining block (left panel), in the near-field area directly adjacent to the mined area (middle), and outside the mining block in the far-field (right panel) under the high disturbance scenario. Immediate impacts are noted in a dark shade and impacts after one year in a lighter shade.

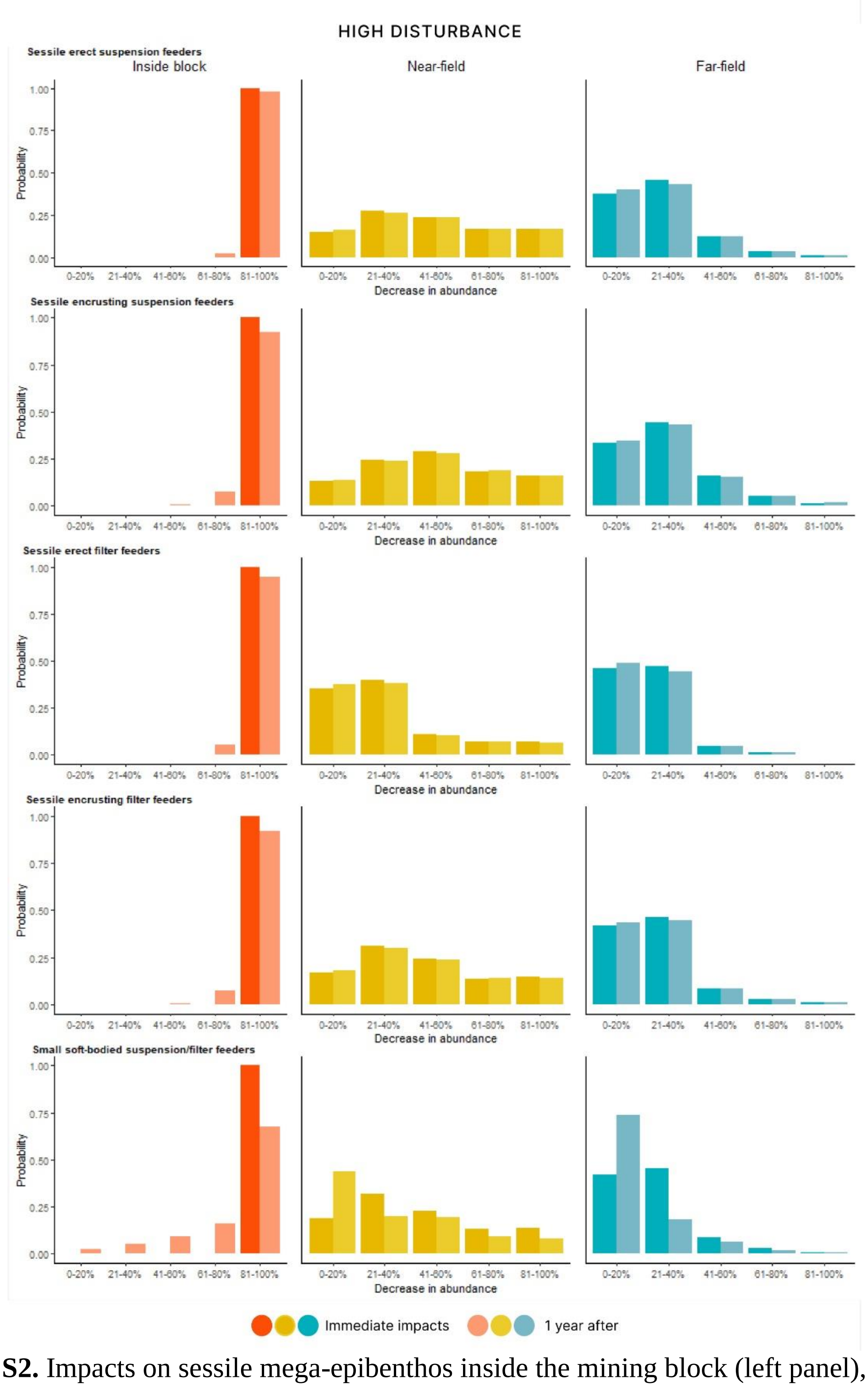


**Figure S2.** Impacts on sessile mega-epibenthos inside the mining block (left panel), in the near-field area directly adjacent to the mined area (middle), and outside the mining block in the far-field (right panel) under the high disturbance scenario. Immediate impacts are noted in a dark shade and impacts after one year in a lighter shade.

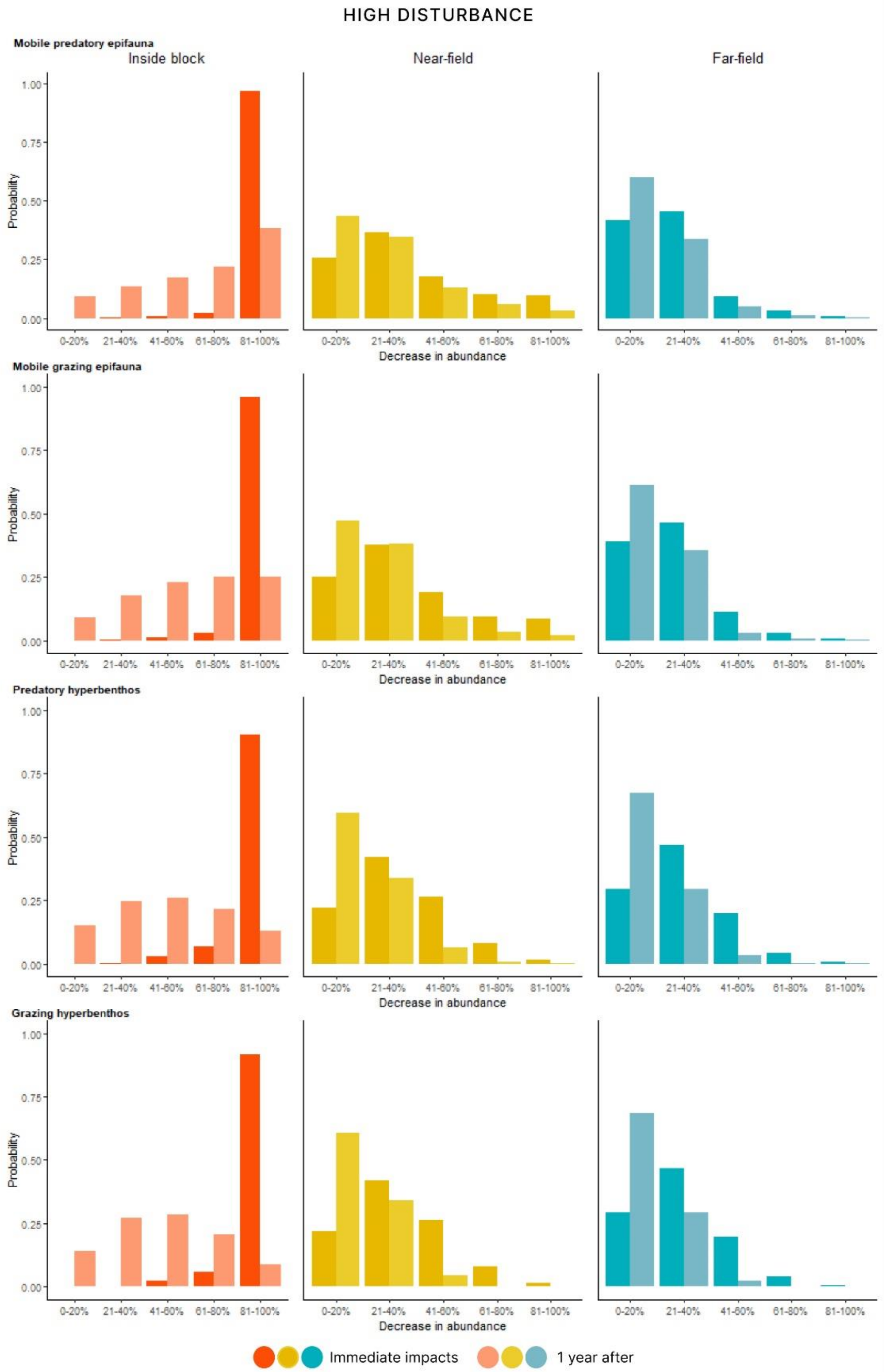


**Figure S3.** Impacts on mobile epibenthos inside the mining block (left panel), in the near-field area directly adjacent to the mined area (middle), and outside the mining block in the far-field (right panel) under the high disturbance scenario. Immediate impacts are noted in a dark shade and impacts after one year in a lighter shade.

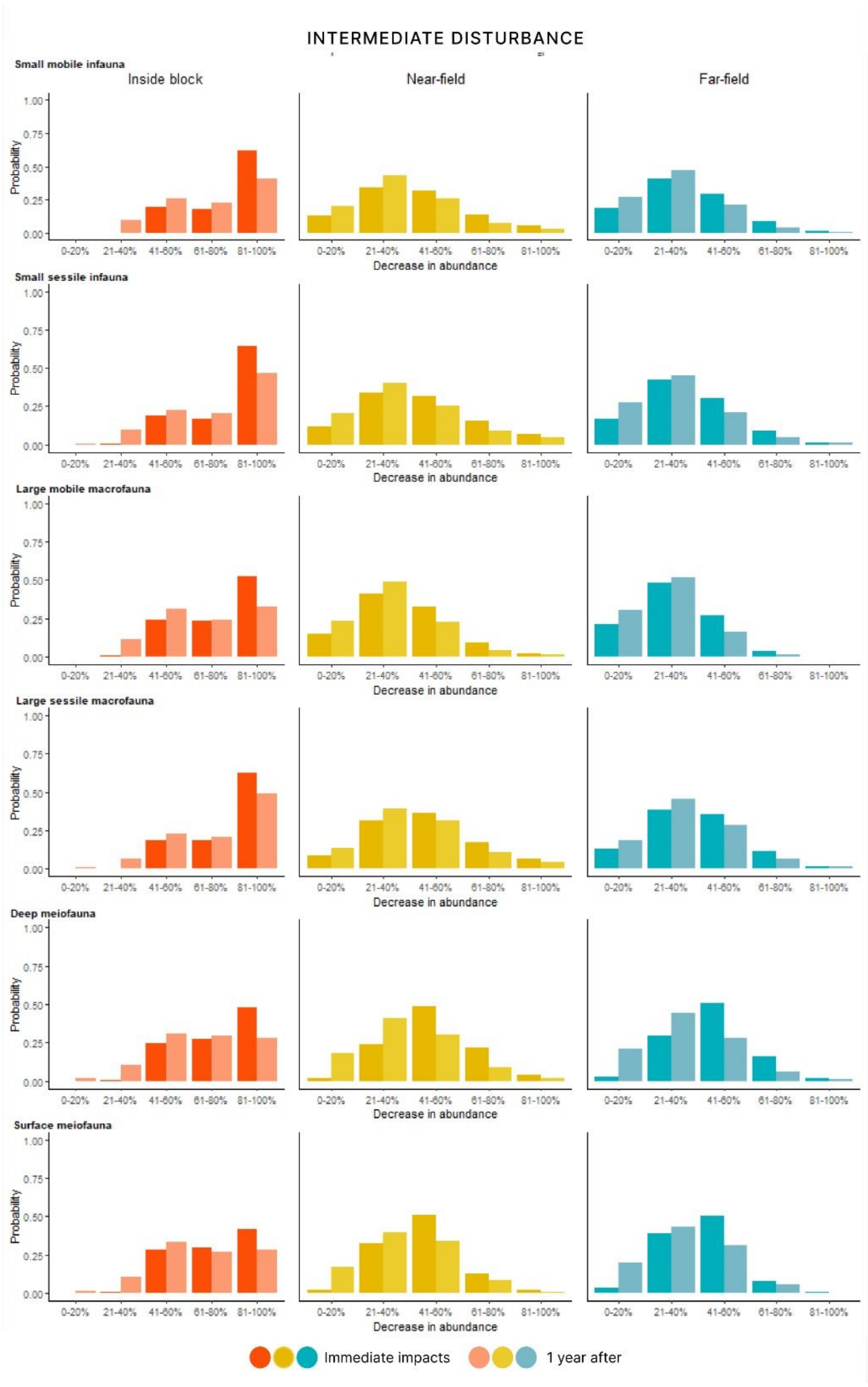


**Figure S4.** Impacts on infauna inside the mining block (left panel), in the near-field area directly adjacent to the mined area (middle), and outside the mining block in the far-field (right panel) under the Intermediate disturbance scenario. Immediate impacts are noted in a dark shade and impacts after one year in a lighter shade.

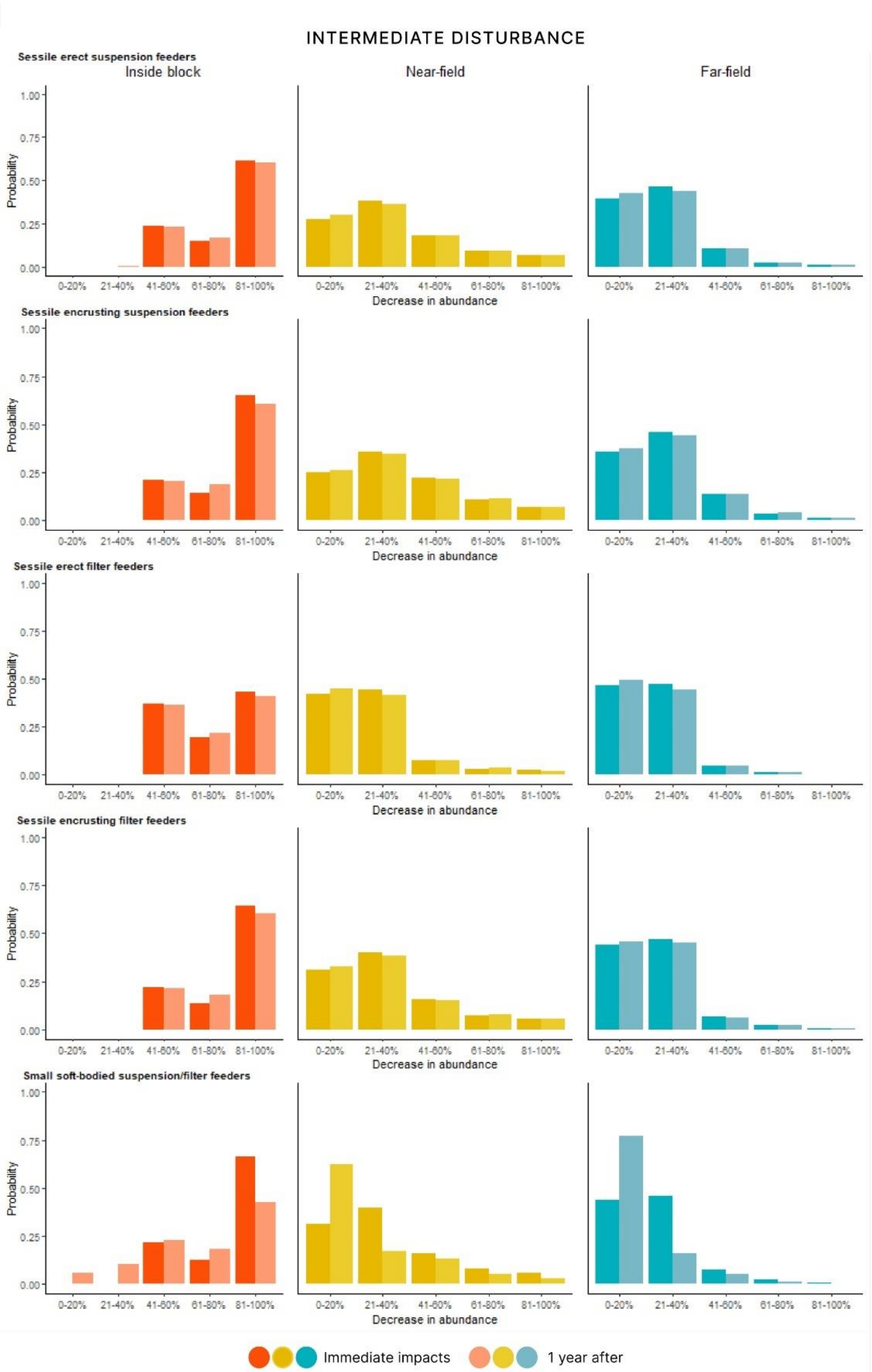


**Figure S5.** Impacts on sessile fauna inside the mining block (left panel), in the near-field area directly adjacent to the mined area (middle), and outside the mining block in the far-field (right panel) under the Intermediate disturbance scenario. Immediate impacts are noted in a dark shade and impacts after one year in a lighter shade.

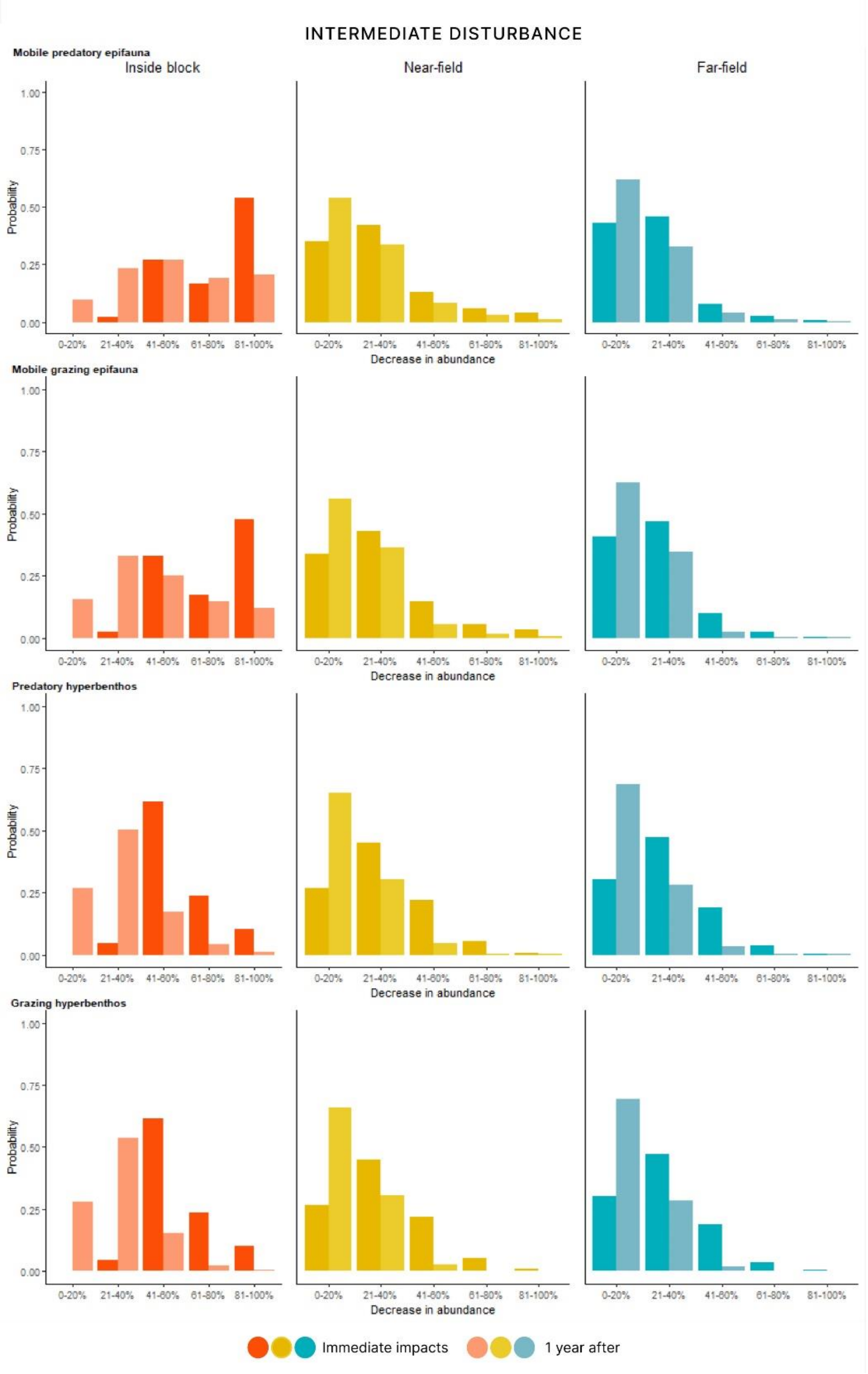


**Figure S6.** Impacts on mobile fauna inside the mining block (left panel), in the near-field area directly adjacent to the mined area (middle), and outside the mining block in the far-field (right panel) under the Intermediate disturbance scenario. Immediate impacts are noted in a dark shade and impacts after one year in a lighter shade.
